# Mast cell activation disrupts interactions between endothelial cells and pericytes during early life allergic asthma

**DOI:** 10.1101/2023.03.07.529253

**Authors:** Régis Joulia, Franz Puttur, Helen Stölting, William J. Traves, Lewis J. Entwistle, Anastasia Voitovich, Minerva Garcia Martín, May Al-Sahaf, Katie Bonner, Elizabeth Scotney, Philip L. Molyneaux, Richard J. Hewitt, Simone A. Walker, Laura Yates, Sejal Saglani, Clare M. Lloyd

**Author notes:** Address correspondence to: Sir Alexander Fleming, Level 3, National Heart and Lung Institute (NHLI), Imperial College London, SW7 2AZ London UK, or Sir Alexander Fleming, Level 3, National Heart and Lung Institute (NHLI), Imperial College London, SW7 2AZ London UK.

## Abstract

Allergic asthma generally starts during early life and is linked to substantial tissue remodelling and lung dysfunction. Although angiogenesis is a feature of the disrupted airway, the impact of allergic asthma on the pulmonary microcirculation during early life is unknown. Here, using quantitative imaging in precision-cut lung slices (PCLS), we report that exposure of neonatal mice to house dust mite (HDM) extract disrupts endothelial cell/pericyte interactions in adventitial areas. Central to the blood vessel structure, the loss of pericyte coverage was driven by mast cell (MCs) proteases, such as tryptase, that can induce pericyte retraction and loss of the critical adhesion molecule N-Cadherin. Furthermore, spatial transcriptomics of paediatric asthmatic endobronchial biopsies suggests intense vascular stress and remodelling linked with increased expression of MC activation pathways in regions enriched in blood vessels. These data provide previously unappreciated insights into the pathophysiology of allergic asthma with potential long-term vascular defects.

## Introduction

Allergic asthma commonly develops during early childhood. While it is difficult to precisely define the prevalence of children affected it is now estimated that between 5% to 20% of all children are impacted by asthma (1, 2). Chronic Inflammation and tissue remodelling are central elements of allergic asthma. Although the immune mechanisms by which allergic responses are well characterised in adult pathologies (3, 4), our understanding of the impact of dysregulated immune responses in early life is only now being appreciated. In addition to airway hyperresponsiveness (AHR) and active tissue remodelling such as increased subepithelial basement membrane thickness, vascular remodelling is also a key feature of allergic asthma (5-7). Indeed, as the disease progresses, the newly altered tissue requires the formation of new blood vessels to efficiently irrigate the remodelled epithelium and adjacent areas (8). The latter has been demonstrated in murine models and human biopsies where an increased number of blood vessels was observed in sub-epithelial areas (9, 10). Furthermore, most of the intermediate to large blood vessels present in the lungs are in close proximity to large epithelial areas described as bronchovascular spaces or adventitia which are involved in multiple pathologies such as asthma and helminth infections (11-14). Central to the maintenance of blood vessels (15-17), little is known about the interaction between endothelial cells and their associated mural cells, pericytes, during lung pathologies (17-19). Pericytes have a unique ability to maintain endothelial cell fitness through the secretion of pro-angiogenic mediators but also cell-cell interaction involving adhesion molecules such as N-cadherin (20).

Whilst lung function starts to decline in children with allergic asthma, the lung vasculature is still developing towards its adult form. Therefore, preventing remodelling during early-life is a critical window of opportunity to prevent disorders later in life. However, the mechanism by which the microcirculation and intermediate blood vessels in close vicinity to large airways is impacted by the responses to allergens remains enigmatic. In this context, mast cells (MCs) are key players in orchestrating the development of allergic asthma through the release of preformed mediators (e.g. histamine, proteases) in response to IgE/antigen or neuropeptide mediated stimulation (3, 21). However, the mechanism by which MCs can regulate vascular inflammation during early life allergic asthma remains to be determined.

Here we used our precision-cut lung slices (PCLS) approach, innovative image quantification, functional assays and spatial transcriptomics to study the lung vasculature in the context of early life allergic airway disease (AAD). Using high dimensional analyses, we discovered that neonatal mice exposed to allergen exhibit vascular remodelling during disease progression, which is defined by a loss of pericyte coverage, reduced density of red blood cells and development of hypoxic areas. Mechanistically, we identified a population of adventitial connective tissue MCs (CTMCs) that were highly activated following allergen exposure. *In vitro*, mouse and human MC granules were able to induce pericyte retraction and reduction of N-cadherin expression. The latter was mediated through MC protease activity and specifically, in human MCs, by tryptase. Finally, spatial transcriptomic data of endobronchial biopsies from children pointed out genetic changes in vessel-rich areas associated with cellular stress, remodelling and MC activation. Together, our data has revealed a new MC/pericyte axis critical in disruption of lung vascular function during early life asthma with potential long-term consequences.

## Results

### Development of a quantitative imaging platform to study lung vasculature in neonatal mice

To determine the impact of early life allergic airways disease on the vasculature, we extended our PCLS approach (22) to simultaneously analyse the different components of the vessel wall, namely: endothelial cells and pericytes. Pericytes are a heterogenous population of cells expressing different phenotypic markers between and within organs (20, 23). We explored the human lung cell atlas data set to identify the most reliable markers of lung pericytes (24). The genes for PDGFRβ and NG2 (*CSPG4*) were the most highly expressed in mouse and human pericytes, and we focussed on PDGFRβ due to its greater level of expression in lung pericytes (Supplemental Figure 1, A-B). PCLS explants obtained from lungs of neonatal mice were stained for CD31 (endothelial cells), α-SMA (smooth muscle cells, SMCs) and PDGFRβ (pericytes) and showed a rich and complex network of blood vessels. Interestingly, pericytes were highly abundant in adventitial regions associated with larger blood vessels but were also present across the parenchyma (Figure 1A, Supplemental Video 1). High magnification images revealed intimate cellular interactions between endothelial cells and pericytes, with extensive protrusions emerging from the pericyte cell body and surrounding endothelial cells (Figure 1B). In addition, PDGFRβ^+^ cells were also positive for NG2 in adventitial and parenchymal areas confirming their pericyte phenotype (Supplemental Figure 1B). We next developed an integrative platform to analyse vascular changes using our PCLS system. This platform relies on the use of tile scan imaging to identify the different lung regions (i.e. bronchovascular space/adventitia and parenchyma) and high-resolution images to define 3D structure and spatial organisation of blood vessels. Using IMARIS software, we processed the images through cell segmentation and volume analyses. Together, we extracted 6 different parameters of vascular inflammation in both the lung adventitia and parenchyma: i) endothelial cell coverage (i.e. volume of endothelial cells within the image), ii) vessel density, iii) pericyte number, iv) pericyte coverage (i.e. volume of pericytes surrounding endothelial cells), v) distance between endothelial cell and pericyte and vi) number of CD45^+^ cells (Figure 1C). Together these parameters provide an accurate representation of the lung vascular architecture and its impact on inflammation.

**Figure 1:**
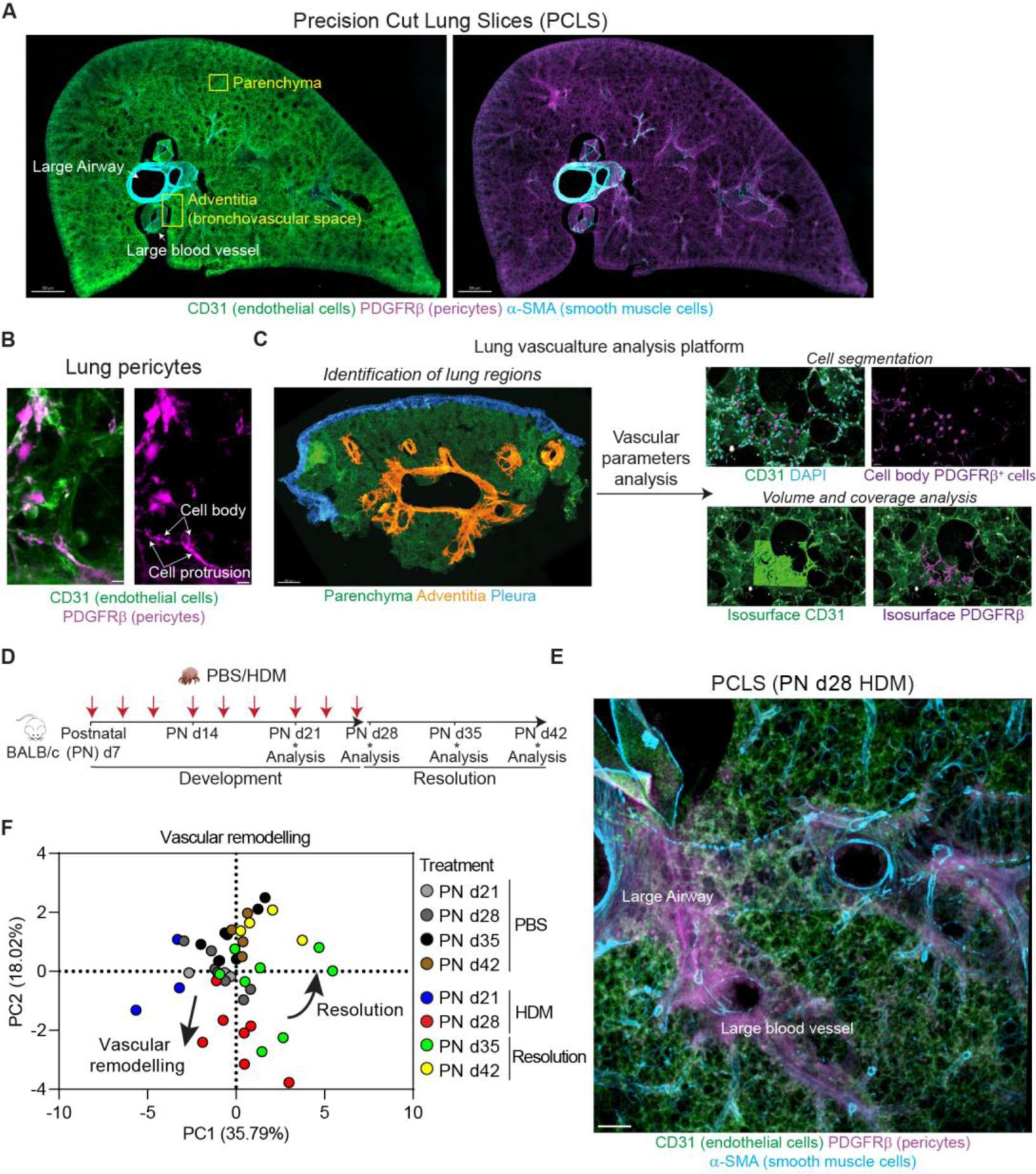
Allergen-induced inflammation leads to vascular remodelling in early life. (**A**) 3D rendering of a PCLS section (200 µm thickness) of neonatal lung (PN d28) stained for CD31 (green, endothelial cells), α-SMA (cyan, SMCs) and PDGFRβ (magenta, pericytes). Yellow box regions indicate adventitial and parenchyma regions analysed (see Supplemental Video 1), scale bars 500 µm (representative of 4 independent experiments). (**B**) Zoomed in image of pericytes (PDGFRβ^+^) extending protrusions around endothelial cells, scale bars 7 µm (representative of 4 independent experiments). (**C**) Image analysis pipeline showing the result of cell segmentation and volume analysis, scale bars 500 µm or 30 µm. (**D**) BALB/c mice aged 7 days were exposed to intermittent intranasal PBS or HDM for 3 weeks (red arrows). Lungs were collected at postnatal (PN) days 21, 28, 35 and 42. (**E**) PCLS section of HDM exposed neonates (PN d28) showing the vasculature in an adventitial region, scale bar 200 µm (representative of 4 independent experiments). (**F**) PCA analysis of lung vascular functions (see Supplemental Figure 2, n= 44 mice from 4 independent experiments).

Next, we employed our platform to analyse the lung vasculature in neonatal mice exposed to HDM (25). Seven-day old BALB/c mice were exposed to intermittent HDM, and lung lobes were collected at different times during disease progression (i.e. postnatal “PN” day 21 and 28) and during the resolution phase after the end of the exposure (i.e. PN day 35 and 42, Figure 1, D-E). As we previously reported for this model, HDM induces all the features of allergic asthma including increased airway resistance, eosinophils, T cells and ILC2 infiltration (25). We performed an unsupervised principal component analysis (PCA) combining the 6 vascular parameters extracted from our images in 2 regions of the lungs (i.e. adventitia and parenchyma). Adventitial regions were identified using structural parameters (i.e. presence of a large airway and intermediate/large blood vessel) and the associated microvasculature within a 150 µm radius from the large airway and large vessel was analysed. Parenchyma was defined using alveolar structures far from any large airway (Figure 1A). Whilst some individual parameters were not significantly different between groups, we did see effects in the total variance when inspecting the principal components of these 12, not independent, parameters. The PCA revealed alterations to the vascular structure as early as 2 weeks following HDM exposure (PN day 21). These changes were further exaggerated by the end of the allergen exposure (PN day 28) and were mainly linked to a loss of the microcirculation. Whilst some mice still exhibited the same vascular changes at PN day 35 during the resolution phase, most had recovered a vasculature comparable to that of age-matched controls at PN day 42 (Figure 1F, loadings, eigenvalues and individual data points shown in Supplemental Figure 2A-K, Figure 3B and Supplemental Figure 4B).

**Figure 2:**
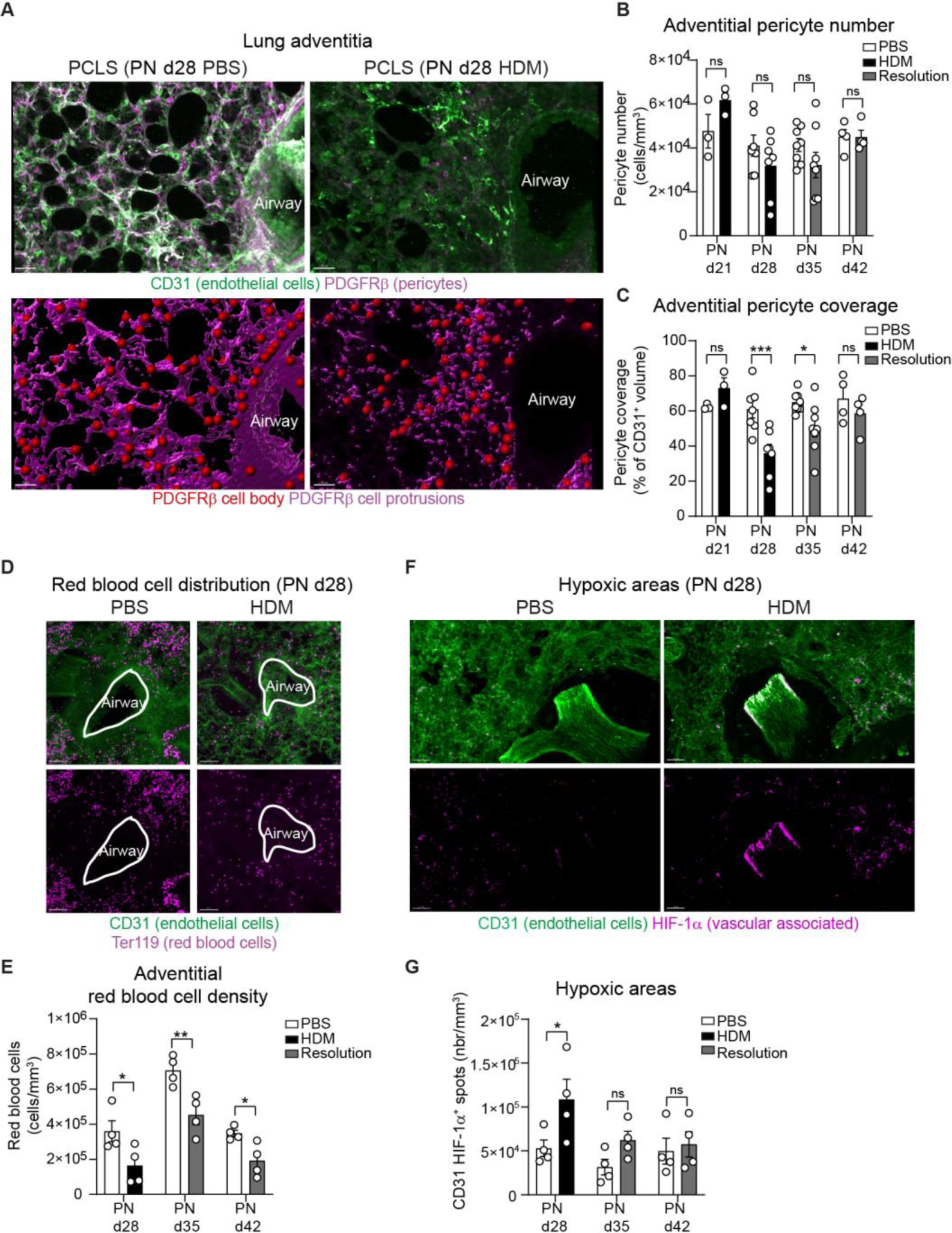
Repeated HDM exposure leads to loss of pericyte protrusions, reduced red blood cells and hypoxic areas. Neonate mice were exposed with PBS or HDM as indicated in Figure 1D. (**A**) 3D rendering of a PCLS section in PBS and HDM exposed mice 3 weeks post first inhalation showing CD31 (green, endothelial cells) and PDGFRβ (magenta, pericytes). Lower panels show the pericyte cell body (red dots) and protrusions (magenta surface) analysis (see Supplemental Video 2), scale bars 30 µm (representative of 4 independent experiments). (**B-C**) PDGFRβ^+^ pericyte number per mm^3^ (B) and coverage (C, normalised to the total volume of CD31^+^ blood vessel) in the lung adventitia (n=3-8 mice per group from 4 independent experiments). (**D**) Representative PCLS showing a reduced red blood cell (Ter119^+^, purple) density in the microcirculation (CD31, green), scale bar 50 µm (representative of 3 independent experiments). (**E**) Adventitial red blood cell density (normalised to the total volume of the image, n=3-4 mice per group from 3 independent experiments). (**F**) PCLS of HDM exposed mice for 3 weeks exhibiting increased HIF-1α (purple) associated to the vasculature (CD31, green), scale bars 30 µm (representative of 4 independent experiments). (**G**) Number of HIF-1α spots in adventitial region in PBS and HDM exposed mice (n= 4 mice per group). Mean ± SEM, two-way ANOVA followed by Sidak’s post-hoc test, *p<0.05, **p<0.01,***p<0.001, ns=not significant.

**Figure 3:**
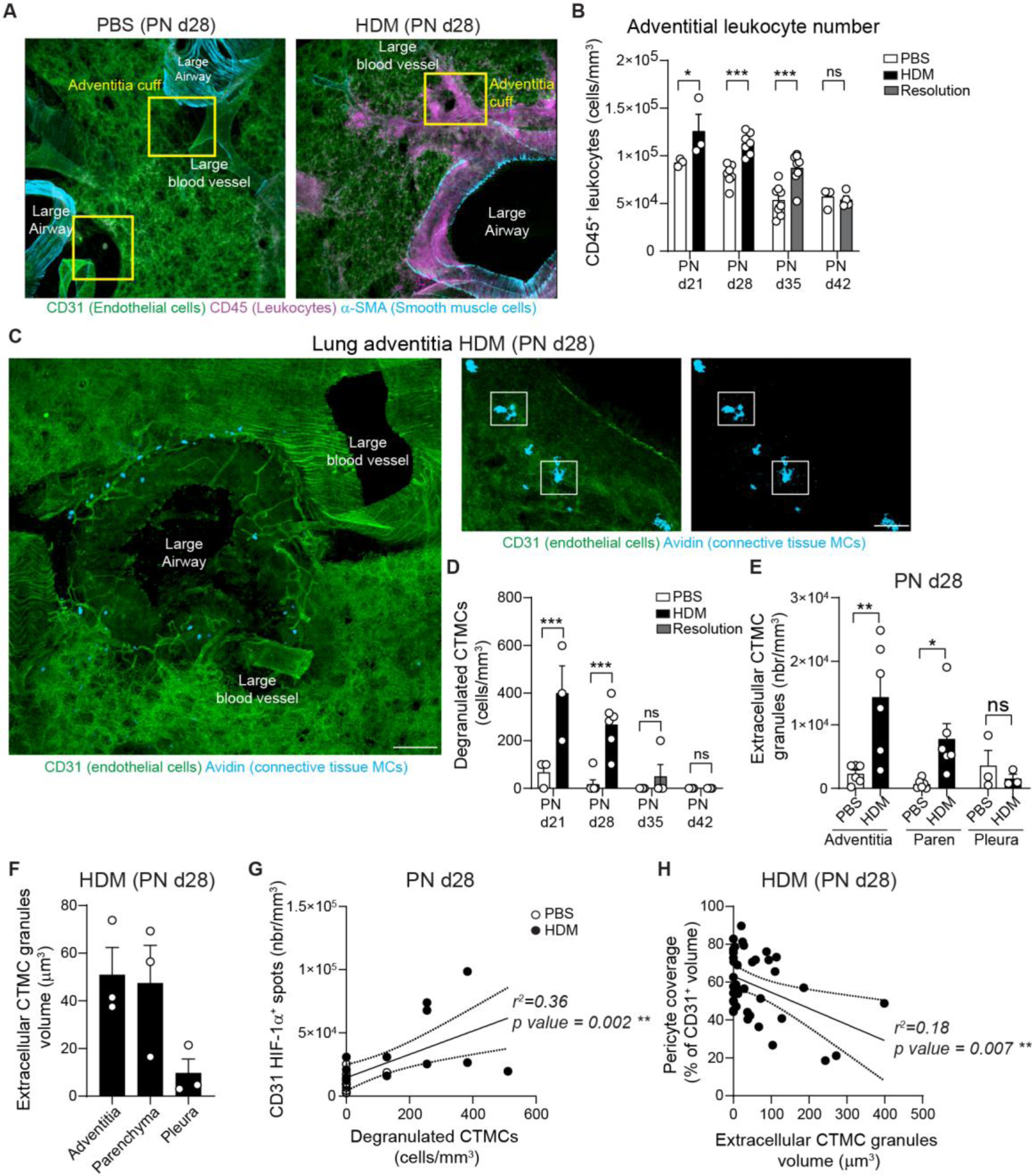
Early life allergen exposure leads to immune cell recruitment and MC activation in the lung adventitia. (**A**) 3D rendering of a PCLS in the lung adventitia in PBS and HDM exposed mice at PN 28 showing CD31 (green, endothelial cells), α-SMA (blue, SMCs) and CD45 (magenta, leukocytes), scale bars 200 µm (representative of 4 independent experiments). (**B**) CD45^+^ cell number (n=3-8 mice per group from 4 independent experiments). (**C**) Representative 3D image of lung adventitia showing the distribution of connective tissue MCs (avidin, blue) around a large airway and associated vasculature (CD31, green) and images showing degranulated MCs adjacent to large airways and blood vessels (see Supplemental Video 3), scale bars 150 µm and 50 µm (representative of 4 independent experiments). (**D**) Number of degranulated MCs (n=3-6 mice per group). (**E-F**) Number of extracellular MC granules per mm^3^ (E) and volume (F) (n=3-6 mice per group from 3 independent experiments). (**G**) Correlation between vascular associated HIF-1α^+^ and number of degranulated MC granules (n=23 images from 4 PBS and 4 HDM treated mice). (**H**) Correlation between pericyte coverage and volume of extracellular MC granules (n=39 images from 4 PBS and 4 HDM treated mice). Mean ± SEM. **B, D, E** two-way ANOVA followed by Sidak’s post-hoc test; **G, H** Spearman’s rank correlation test. *p<0.05, **p<0.01,***p<0.001, ns=not significant.

In summary, our data clearly indicate that HDM exposure induced vascular remodelling in early life mainly due to vascular loss. These changes are maintained during the resolution phase but slowly resolve with time.

### Loss of pericyte coverage occurs early in HDM exposed neonatal mice

One of the principal parameters determining the vascular remodelling in our PCA analysis (Figure 1F) was the change in pericyte coverage in close proximity to the lung adventitia. Pericyte coverage, or the volume of pericyte cell protrusions, is an essential factor to maintain endothelial cell fitness and overall blood vessel structure such as in the brain where they form the blood brain barrier (15). During early life AAD, zoomed in images clearly indicate that whilst the endothelial cell structure was still present in close vicinity to large airways at PN day 28, pericyte signal was greatly reduced (Figure 2A & Supplemental Video 2). Interestingly, our image analysis revealed that pericyte cell bodies were still present in PBS and HDM groups but the extent of pericyte protrusions was severely reduced following 3 weeks HDM exposure (Figure. 2A). Indeed, quantitative analysis confirmed that adventitial pericyte cell number was not significantly changed during mouse development or impacted by HDM exposure (Figure 2B). In contrast, pericyte coverage was reduced after 3 weeks of allergen exposure (i.e. ∼42% reduction) and slowly recovered in the resolution phase to nearly physiological levels 2 weeks post last HDM inhalation (Figure 2C). Endothelial cell volume or blood vessel density did not show strong differences however we detected a small reduction in endothelial cell volume at the end of the HDM exposure and this decrease was significant 1 week into the resolution phase (Supplemental Figure 2E). The latter may indicate that the reduction in pericyte coverage leads to a decrease in the microcirculation of lung adventitial regions. Finally, we assessed the functional consequences of the adventitial vascular changes. To this end, we analysed the distribution of red blood cells and the expression of hypoxia-inducible factor 1α (HIF-1α) in the vasculature (i.e. CD31^+^ areas). Following 3 weeks of HDM exposure, the adventitial vasculature showed reduced red blood cell density (Figure 2, D-E) and increased expression of HIF-1α (Figure 2, F-G). Interestingly, whilst HIF-1α slowly came back to normal in the resolution phase (Figure 2G), the red blood cell density reduction was still present 2 weeks post end of challenge (Figure 2E). To investigate the long-term consequences of early life AAD, we treated neonates for 3 weeks with HDM and re-exposed mice to a single dose of allergen or PBS at PN day 42; 2 weeks after last HDM exposure (Supplemental Figure 3A). Whilst the mice re-treated with HDM did not exhibit a reduction in pericyte number compared to control animals, pericyte coverage was significantly reduced (∼26% reduction, Supplemental Figure 3B-D). In addition, we did not observe changes in other vascular parameters in the lung adventitia (i.e. distance endothelial cell/pericyte, endothelial cell volume and blood vessel density) but an increased expression of HIF-1α indicating local change in oxygen concentration (Supplemental Figure 3E-I). These data clearly indicate that allergen exposure during early life leads to functional consequences on the vasculature that can be long lasting.

Collectively, these data demonstrate, for the first time, that repeated exposure to allergen during the early life developmental period induces loss of pulmonary pericyte protrusions and blood vessels leading to hypoxic areas in the pulmonary microcirculation.

### Lung adventitia is marked by immune cell infiltration and mast cell activation

Lung adventitial regions are characterised by an abundant presence of resident immune cells coupled with intense recruitment of various leukocytes during inflammation (14, 22, 26). We hypothesized that local activation of the immune system may play a role in the vascular remodelling and changes in pericyte morphology observed in our neonatal mice following inhaled HDM exposure. Therefore, we employed our image analysis platform to dissect immune cell recruitment and activation following HDM exposure (Figure 3A) in the lung adventitia and parenchyma. We observed a significant increase in the number of adventitial CD45^+^ leukocytes following each HDM exposure (i.e. ∼36% and ∼40% increase after 2 and 3 weeks of HDM, respectively), with the difference still present 1 week into the resolution phase (i.e. ∼60% increased), and returning to physiological levels 2 weeks following the end of the allergen exposure period (Figure 3, A-B). Immune cell recruitment was mainly restricted to the periadventitial regions since the lung parenchyma did not show a significant increase in leukocytes (Supplemental Figure 4, A-B).

Given that MCs play a key role in allergic asthma, and we recently demonstrated the impact of perivascular connective tissue mast cells (CTMCs) in regulating pericyte function (27), we hypothesized that MCs could play a role in mediating pericyte change during early life AAD. We utilised fluorescent avidin staining to analyse the profile of degranulated CTMCs as we and others have demonstrated that it provides an accurate measurement of the localisation and cellular interactions of extracellular CTMC granules (28-31). Whilst the number of CTMC did not change in HDM exposed neonatal mice (Supplemental Figure 4C), we observed a clear increase in the number of degranulated CTMCs every time mice were challenged with allergen (Figure 3, C-D and Supplemental Video 3). We found extracellular CTMC granules were abundant in the lung adventitia and within the parenchyma but not in the lung pleural cavity (Figure 3E). These granules retain a large volume (i.e. ∼50 µm^3^) even outside of the MC cell body and no differences were observed between adventitial and parenchymal MC granule volume (Figure 3F, Supplemental Video 3). Interestingly, we observed a positive correlation between the extent of degranulated CTMCs and expression of HIF-1α (r^2^= 0.36, p value= 0.002, Figure 3G). Furthermore, the volume of CTMC granules was negatively correlated with the reduction in pericyte coverage (r^2^= 0.18, p value= 0.007, Figure 3H) indicating that regions with large extracellular CTMC granules are associated with change in pericyte morphology. Finally, the presence of CTMC granules was linked with an increased distance between endothelial cells and pericytes (Supplemental Figure 4D).

In summary, our data show that upon exposure to allergen, neonatal mice develop strong adventitial inflammation linked with activation and degranulation of resident CTMCs. Areas with large CTMC granules are associated with destabilisation of endothelial cell/pericyte interaction that likely disrupt the vasculature leading to hypoxic areas.

### CTMC degranulation induces pericyte retraction and loss of N-Cadherin

To analyse the functional impact of CTMC degranulation on lung pericytes, we developed an *in vitro* co-culture model between primary lung MCs and pericytes. Lung cells were purified from naïve mouse lungs and following 2-4 weeks in culture, a high level of purity was detected for both cell populations with expression of characteristic markers: NG2^+^/PDGFRβ^+^ and FcεRI^+^/ST2^+^/CD117^+^ for pericytes and MCs respectively (Supplemental Figure 5A). In addition, lung MCs were positive for avidin signal indicating that the *in vitro* generated MCs were indeed CTMCs (Supplemental Figure 5A). Anti-dinitrophenyl (DNP) IgE sensitized lung MCs were added to a layer of fluorescently labelled pericytes (CMTMR^+^) and stimulated with increasing concentrations of DPN-BSA to induce FcεRI crosslinking and degranulation (Figure 4A). Degranulated lung MCs were identified using extracellular avidin staining revealing the presence of exteriorised granules in the surrounding environment or still associated with the surface of MCs (Figure 4B). Whilst the frequency of degranulated MCs increased proportionally with the concentration of DNP-BSA (Figure 4C), the number of pericytes was not impacted by MC activation status (Figure 4B and D). However, as observed in vivo, the volume of pericytes decreased significantly 24h after stimulation (∼30% reduction for 100 ng/ml DNP-BSA, Figure 4B and E). We next investigated if the reduction in volume was more pronounced in pericytes that were directly in contact with MC granules. Avidin mean fluorescence intensity (MFI) on the largest pericytes (i.e. >45000 µm^3^) showed lower avidin signal compared to the smallest pericytes (i.e. <17000 µm^3^) indicating that the smallest pericytes had more MC granules on their surface that could induce their retraction (Figure 4F). Furthermore, one of the main consequences of pericyte retraction is the polarisation of F-actin (32). Indeed, concomitant with reduced pericyte volume, the increased number of degranulated MCs led to an enhanced intracellular F-actin signal per unit of volume within pericytes (Figure 4, G-H). Interestingly, N-Cadherin expression, one of the main cadherins involved in the interaction between endothelial cells and pericytes (33) was decreased following contact with MC granules (∼20% reduction at 100 ng/ml DNP-BSA, Figure 4I).

**Figure 4:**
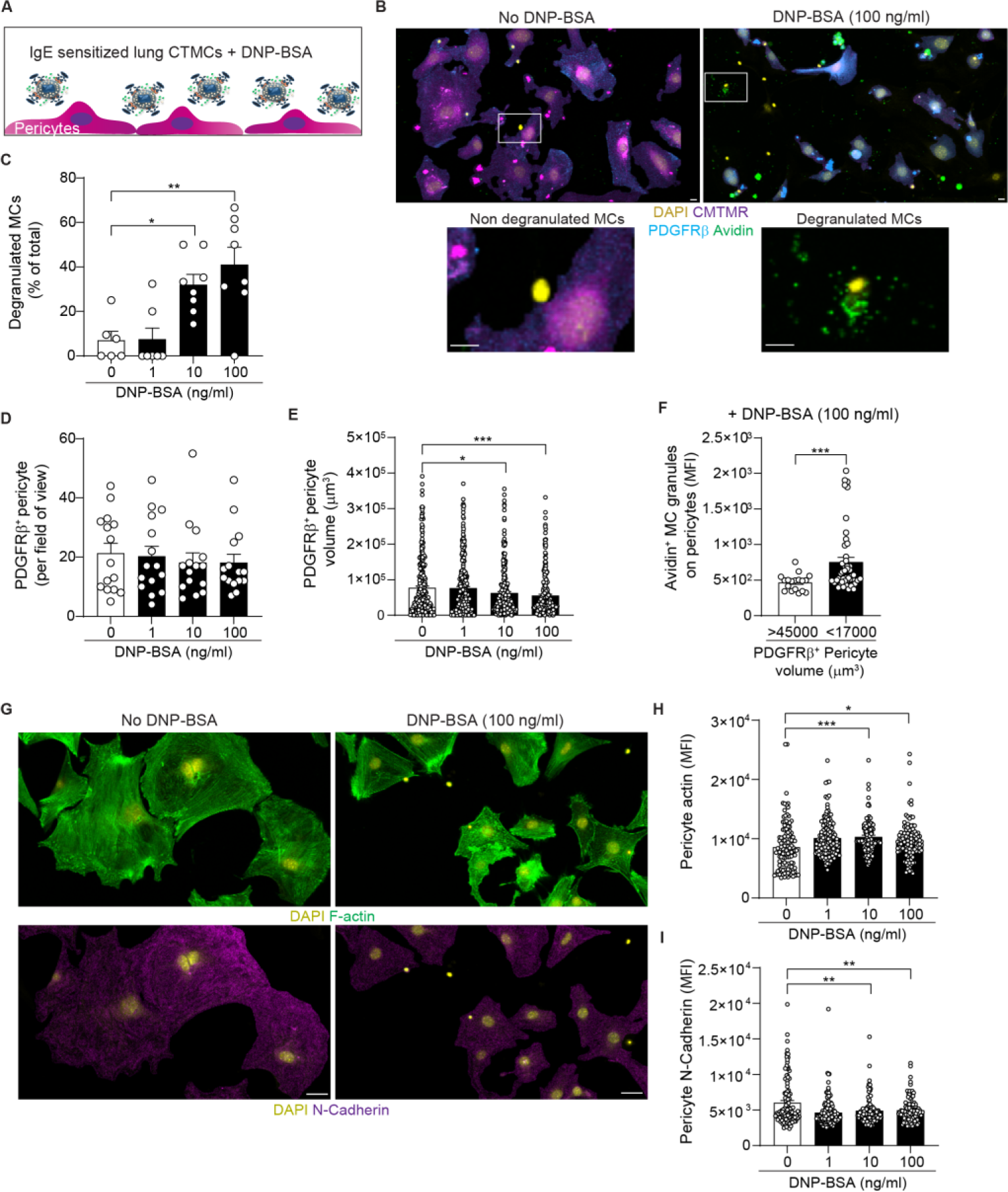
MC granules induce pericyte retraction and cleavage of surface N-cadherin. MCs were sensitized overnight with anti-DNP IgE then placed on a layer of pericytes (stained with CMTMR) and stimulated with increasing concentrations of DNP-BSA for 24 hours. (**A**) Schematic depicting the coculture experiment between primary mouse lung MCs and pericytes. (**B**) Images of unstimulated or stimulated (100 ng/ml DNP-BSA) pericyte/MC co-cultures stained for DAPI (yellow), CMTMR (purple), PDGFRβ (blue) and avidin (green, MC granules), white boxes indicate the areas zoomed in showing examples of resting or degranulated MCs, scale bars 15 µm (representative of 3 independent experiments). (**C**) Frequency of degranulated MCs (n=6-8 images from 3 independent experiments). (**D**) Number of pericyte per field of view (n=15 images from 3 independent experiments). (**E**) Pericyte volume determined using the cell tracer CMTMR (n=253-299 pericytes from 3 independent experiments). (**F**) Avidin signal on small pericytes (<17000 µm^3^) and large pericytes (>45000 µm^3^) showing that small pericytes exhibit more MC granule staining on their surface (n=18-46 from 3 independent experiments). (**G**) Representative images of F-actin (green) and N-Cadherin (magenta) pericyte expression in the presence of degranulated MCs or control, scale bars 50 µm (representative of 3 independent experiments). (**H-I**) F-Actin (H, n=98-144) and surface N-Cadherin (I, n=98-144) MFI on pericytes (3 independent experiments). Mean ± SEM. **C**, **D**, **E**, **H**, **I** one-way ANOVA followed by Tukey’s post-hoc test; **F**, two-tailed Student’s t-test. *p<0.05, **p<0.01, ***p<0.001, ns=not significant.

Collectively, MC granules can efficiently induce pericyte retraction in a dose-dependent manner and facilitate reduction in N-Cadherin expression.

### Pericyte retraction and N-Cadherin loss are mediated by MC proteases

Having observed a reduced pericyte volume and N-Cadherin expression on pericytes following MC degranulation, we next investigated the molecular mechanisms of MC induced pericyte retraction. MCs contain a vast repertoire of proteases, which are involved in protective immunity against venoms and parasites but can also be detrimental in the context of inflammatory pathologies such as asthma (34, 35). We hypothesized that MC derived proteases could be involved in pericyte retraction. Indeed, PCLS of HDM exposed neonates (PN d28) indicated that proteases such as m-MCP6 (MC tryptase) were present in CTMC granules (Figure 5A). Furthermore, detailed colocalization analysis indicated the presence of m-MCP6 in both intracellular and extracellular MC granules, 24 hours post last HDM challenge, as demonstrated by their overlap with avidin^+^ regions (Figure 5B). The latter agrees with the requirement for mMCP-6 to be associated with the CTMC granule matrix to be stored and activated efficiently (35, 36). We next tested if a general protease inhibitor could prevent pericyte protrusions loss in the presence of degranulated MCs. Interestingly, our results showed that although MC degranulation was not impacted by the protease inhibitor (Figure 5, C-D), pericyte volume and N-Cadherin surface expression were maintained (Figure 5, E-F). To further confirm this observation, we directly exposed lung pericytes to recombinant m-MCP6 and observed a ∼62% reduction in pericyte volume indicating that m-MCP6 at a high concentration is sufficient to induce pericyte retraction (Figure 5G).

**Figure 5:**
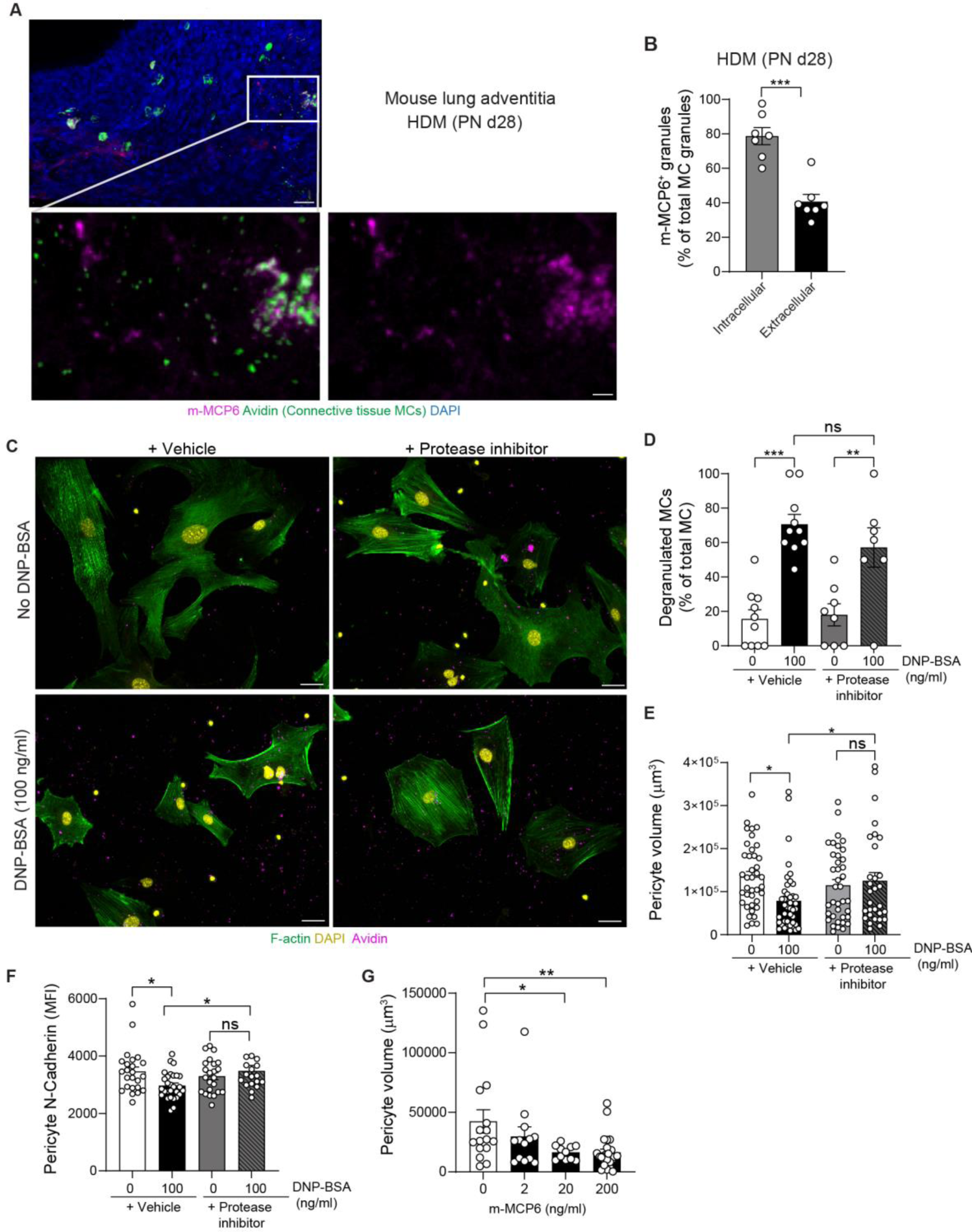
MC derived proteases induce pericyte retraction and N-Cadherin cleavage. (**A**-**B**) Neonate mice were exposed to HDM for 3 weeks. (**A**) 3D rendering of a PCLS section in the lung adventitia in HDM exposed mice showing DAPI (blue), m-MCP6 (mouse tryptase, magenta) and MCs (avidin, green), lower panels show zoomed-in images of the white box region and the m-MCP6 signal in extracellular MC granules, scale bars 30 and 10 µm. (**B**) Colocalization analysis between intracellular and extracellular MC granules (avidin^+^) and m-MCP6 showing the frequency of m-MCP6^+^ granules (each dot represents an image from 3 independent mice). (**C**-**F**) MCs were sensitized overnight with anti-DNP IgE then placed on a layer of pericytes (stained with CMTMR) and stimulated with increasing concentration of DNP-BSA for 24h in the presence of a protease inhibitor cocktail or vehicle control (DMSO). (**C**) Images of unstimulated or stimulated (100 ng/ml DNP-BSA) pericyte/MC co-cultures stained for DAPI (yellow), F-actin (green) and MC granules (avidin, magenta), scale bars 50 µm (representative of 3 independent experiments). (**D**) Number of degranulated MCs (normalised to the total number of MCs, each dot represents an image from 3 independent experiments). (**E**) Pericyte volume and (**F**) cell surface N-cadherin MFI on pericytes (each dot represents an individual pericytes from 3 independent experiments). (**G**) Lung pericyte volume 24h following recombinant m-MCP6 exposure (each dot represents an individual pericyte from 3 independent donors). Mean ± SEM. **B**, two-tailed Student’s t-test; **D**, **E**, **F**, **G** one-way ANOVA followed by Tukey’s post-hoc test *p<0.05, **p<0.01, ***p<0.001, ns=not significant.

Together, these data demonstrate that MC proteases can efficiently induce pericyte retraction and loss of surface N-cadherin and potentially lead to loss of the endothelial cell/pericyte interaction.

### Spatial transcriptomic analysis suggested cellular stress in vessel rich areas of children with asthma and human lung pericytes retract following MC degranulation

We next employed digital spatial profiling with the NanoString GeoMx^®^ Cancer Transcriptome Atlas (CTA, ∼1,800 genes) to analyse vascular remodelling and MC activation in 4 children with severe asthma (between 9 and 17 years old, Supplemental Table 1) and 2 controls (between 8 and 11 years old, Supplemental Table 1). Formalin-fixed paraffin-embedded (FFPE) sections obtained from endobronchial biopsies were stained for DNA, Vimentin, CD45 and α-SMA to determine 4 types of regions namely: epithelium (defined using morphology and DNA stain), immune cell infiltrate (CD45^+^ rich), smooth muscle, and fibroblast-rich areas (Figure 6A) (37).

**Figure 6:**
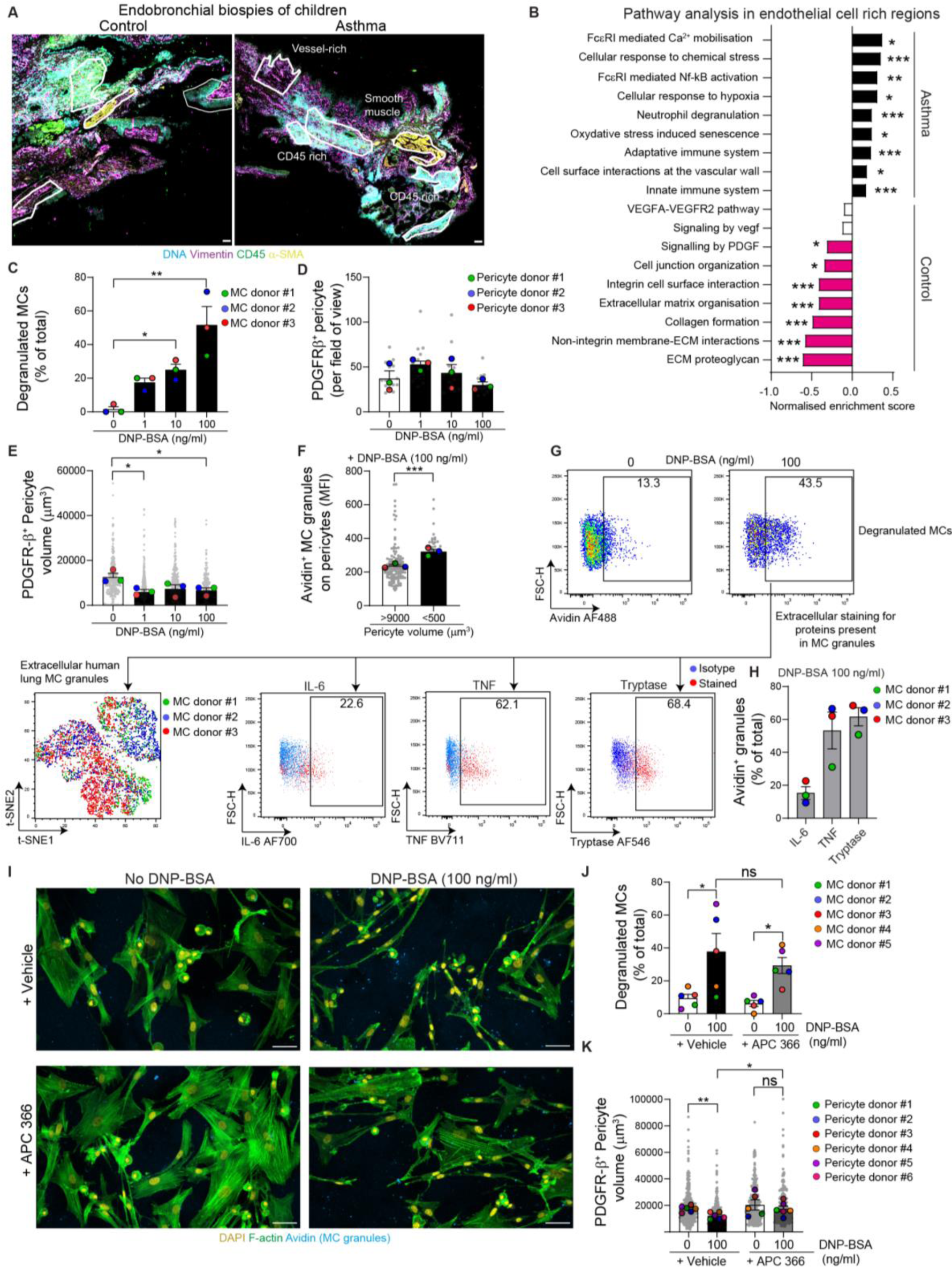
Transcriptional signature of children with asthma suggests vascular stress and human MCs induce tryptase dependant pericyte retraction. (**A**) Immunofluorescence images of endobronchial biopsies stained for DNA (Syto83, green), Vimentin (purple), CD45 (blue) and α-SMA (yellow) showing the selected ROIs (boxed regions), scale bars 100µm. (**D**) Pathway enrichment analysis in endothelial cell rich regions of children with asthma and controls (n=4-9 ROIs per group from 2 controls and 2 asthma, p values in Supplemental Table S4). (**C-F**) and (**I-K**), Human MCs were sensitized overnight with anti-DNP IgE then placed on pericytes and stimulated with an increasing concentration of DNP-BSA for 24h, (**I-K**) APC366 (tryptase inhibitor) or vehicle control were added at the time of stimulation. (**C**) Degranulated MCs number (normalised to the total number of MCs, n=3 MC donors from 3 independent experiments). (**D**) Pericyte number (n=3 Pericyte donors from 3 independent experiments). (**E**) Pericyte volume (n=3 Pericyte donors from 3 independent experiments). (**F**) Avidin signal on small pericytes (<500 µm^3^) and large pericytes (>9000 µm^3^) (n=3 Pericyte donors from 3 independent experiments). (**G**) Flow cytometry profiles of degranulated MCs and unsupervised analysis of extracellular MC granules (t-SNE analysis performed on 3 pooled donors, representative of 2 independent experiments). (**H**) Frequency of avidin^+^ granules positive for indicated markers (n=3 MC donors from 2 independent experiments). (**I**) Images of pericyte/MC co-culture stained for DAPI (yellow), F-actin (green) and MC granules (avidin, blue), scale bars 50µm (representative of 3 independent experiments). (**J**) Degranulated MCs number (normalised to the total number of MCs, n=5 MC donors from 3 independent experiments). (**K**) Pericyte volume (n=6 pericyte donors from 3 independent experiments). Mean ± SEM. **B**, two-tailed Mann Whitney test; **C**,**D**,**E**, one-way ANOVA followed by Tukey’s post-hoc test; **F**, two-tailed Student’s t-test; **J**,**K**, two-way ANOVA followed by Sidak’s post-hoc test. *p<0.05, **p<0.01,***p<0.001, ns=not significant.

We analysed 40 ROIs (Supplemental Table 2) and first defined the regions with the highest density of blood vessels using endothelial cell and pericyte cell deconvolution using single cell RNA-seq cell-type signatures (38) (genes employed are indicated in Supplemental Table 3). We selected 16 ROIs from 2 controls and 2 patients with asthma with high endothelial cell signature (i.e. enrichment score >0.15) in fibroblast and immune cell infiltrate areas (Supplemental Figure 6A). Whilst the pericyte enrichment score was not statistically different across the 4 types of regions (Supplemental Figure 6B), we observed a correlation between regions with high endothelial cell and pericyte signatures, suggesting that these regions are enriched in microvessels (Supplemental Figure 6C). The high endothelial cell signature was further suggested by the higher expression of key vascular genes (i.e. *CDH5*, *PECAM1*, *PDGFRB*) in the 16 ROIs identified (Supplemental Figure 6D). Gene set enrichment analysis (GSEA) was employed to analyse pathways differentially regulated between control and asthma ROIs in endothelial cell rich regions between controls. Although this analysis is limited by the number of ROIs available, patients with asthma had increased signals in pathways related to cellular stress and hypoxia and downregulation of pathways linked to extracellular matrix organisation and PDGF signalling (Figure 6B, uncorrected and corrected p values are indicated in Supplemental Table S4). Individual gene values for critical pathways (e.g. cellular response to chemical stress, cellular response to hypoxia, signalling by PDGF and extracellular matrix organisation) are shown in Supplemental Figure 6E. Of note, this preliminary analysis suggested no differences in endothelial cell and pericyte enrichment between control and asthma ROIs (Supplemental Figure 6F). Interestingly, MC activation genes such as FcεRI activation pathway were particularly enriched in endothelial cell regions of patients with asthma (Figure 6B). Cell deconvolution analysis hinted a high proportion of MCs in fibroblast and CD45 rich areas (Supplemental Figure 6G) that correlated with the endothelial cell enrichment score (Supplemental Figure 6H). The increased proportion of MCs in endothelial cell rich regions was further suggested by an increase in expression of the tryptase gene (i.e *TPSAB1*, Supplemental Figure 6I) and no difference in abundance were observed between controls and asthma in endothelial cell rich regions (Supplemental Figure 6J).

Finally, we isolated human pericytes and MCs from healthy lung tissue and performed co-culture experiments. Isolation and culture methods yielded purities of more than 95% for both cell types (Supplemental Figure 5B). We exposed lung pericytes to IgE-sensitized MCs and increasing concentrations of DNP-BSA for 24h leading to MC degranulation without impacting pericyte viability (Figure 6, C-D). As observed for murine cells, human pericytes retract following MC activation (Figure 6E) and the volume reduction was enhanced in cells in close contact with MC granules (Figure 6F). To get a better insight into the molecular mechanism of MC dependent pericyte retraction, we analysed the content of extracellular MC granules by flow cytometry. Following 30 min IgE-mediated stimulation, we performed an unsupervised analysis (i.e. t-SNE) of membrane-bound MC granules (i.e. avidin^+^ MC (31)) pooling 3 independent stimulated lung MC donors (Figure 6G). We discovered strong heterogeneity in the profile of extracellular MC granules with the presence of cytokines such as IL-6 and TNF and an abundant presence of tryptase in ∼61.6% of extracellular MC granules (Figure 6H). Since tryptase is highly expressed in extracellular MC granules, we hypothesized that this protease could directly induce pericyte retraction. Therefore, we performed co-culture experiments between activated MCs and pericytes in the presence of a specific tryptase inhibitor (i.e. APC-366) (31, 39). Whilst APC-366 did not impact MC degranulation (Figure 6, I-J), pericyte volume was maintained in the presence of the inhibitor (Figure 6I and K).

Overall, our data indicate that tryptase released from human MCs induces loss of pericyte protrusions.

## Discussion

Allergic airway diseases disproportionally impact children at a stage of their life when the lungs are still developing structurally - including the vast vascular network. Despite our growing understanding of the distinctions between adult and neonatal immune responses (25, 40, 41), the impact of respiratory diseases such as allergic asthma on the lung vasculature remains elusive. Tissue remodelling such as increased thickness of the reticular basement membrane and collagen deposition is a key feature of chronic inflammation and angiogenesis is classically associated with this phenomenon (7, 25, 42). Indeed, as the tissue adjacent to the airway is expanding, newly developed bronchial blood vessels are required to supply the necessary nutrients. Paradoxically, how the original pulmonary microcirculation in adventitial regions adapt to the newly developed tissue currently remains unclear. Here, we demonstrate that pulmonary exposure to inhaled allergens such as HDM during early life drives important vascular changes in the lung adventitia and bronchovascular space. For the first time, we provide an accurate characterisation of the cascade of events leading to vascular remodelling starting with a reduction in pericyte coverage, associated with reduced red blood cell density concomitant with increased hypoxic regions in specific lung regions. We show that the loss of protrusions from the pericyte surface is driven by proteases released following mast cell degranulation. Finally, spatial transcriptomics analysis pointed out a strong remodelling signature in vessel-rich areas of lungs from children with asthma associated with increased MC activation that can induce human pericyte retraction.

Endothelial cell-pericyte interaction is an essential component of the stability of blood vessels however the mechanisms by which this interaction is dysregulated during inflammation remain poorly understood (43). Here, we identified that whilst the number of pericytes did not change dramatically throughout post-natal lung development or following exposure to allergens, the reduction in pericyte coverage was one of the earliest events observed for the vasculature during the development of AAD. The role of pericytes in lung biology is still poorly understood, fate mapping and depletion models provided the first evidence for the involvement of pericytes in lung pathologies such as pulmonary hypertension and lung fibrosis (44, 45). Our data concur with previous work showing that in adult mice, the disruption of the PDGFβ-PDGFRβ axis worsened lung function such as airway hyperreactivity (18). The reduction in pericyte coverage is reminiscent of other pathologies such as Alzheimer’s disease or following stroke. The latter drives a reduction in blood flow in specific areas leading to a decline in brain function (46, 47). Mechanistically, we show that mouse and human MC-derived granules induce pericyte retraction. Pericytes respond to a variety of inflammatory stimuli such as ROS, prostaglandin E2 and TNF (43) however, to our knowledge, the impact of MCs on pericyte coverage has never been described before. We identified that MC-derived proteases such as tryptase induce pericyte retraction and loss of N-Cadherin. The latter is a crucial junctional molecule mediating endothelial cell-pericyte interactions (48) and further studies are required to understand how MC proteases can cleave N-cadherin. In addition, other proteases such as neutrophil derived proteases present in neutrophil extracellular traps (NETs) (49) may play a role in pericyte damage that remain to be investigated. The physiological relevance and mechanism by which pericyte retraction occurs are still not fully elucidated. The nature of this phenomenon is difficult to analyse in vivo, and in vitro experiments may not accurately represent how pericytes contract in vivo. Recent developments to provide better markers for pericytes (50) and improved in vitro models will enhance our understanding of this enigmatic cell.

Lung adventitial regions have recently received great interest and spatial analysis has started to uncover their specialised immune regulation (13, 14). In this context, we showed that lung adventitial regions were central to the inflammatory and remodelling response during early life AAD. Indeed, most of the vascular changes and inflammatory cell recruitment observed were confined to this region, confirming the essential role of spatial approaches to analyse lung immune responses. However, the limit of the lung adventitia, identity of pericytes in these regions and whether there is a subclass of pericytes operating during development that are more susceptible to inflammation remains to be determined (51). Furthermore, the complexity of human airways is not perfectly reflected in murine models and additional work is necessary to provide a better characterisation of human adventitial areas.

We identified a population of connective tissue MCs present around the airway and large blood vessels in the neonatal developing lung. This agrees with previous studies which showed an increase in MC numbers from day 7 after birth (40). Our data extend this observation by determining the cellular location of MCs and explores their functional involvement during AAD. The specific distribution of MCs in these regions during early life remain unclear but the presence of mural cells such as vascular smooth muscle or subtype of pericytes could lead to the development of MCs (52). In addition, IL-33 is a potent MC activator able to regulate the extent of MC degranulation and production of chemokines (53). However, despite evidence that the mechanisms of release of IL-33 between mouse and human is different, it is established that IL-33 is critical for immune responses during early life (42, 54). The precise impact of IL-33 on MC activation during early life remains to be explored. Here, we employed an IgE-mediated MC stimulation to analyse the impact of MC proteases on pericytes in vitro. Whilst we cannot rule the involvement of other MC stimuli such as substance P, HDM treated neonatal mice mount efficient IgE responses following 3 weeks of allergen exposure (54, 55) thus validating our in vitro approach.

Interestingly, our preliminary spatial transcriptomic analysis of endobronchial biopsies from children with asthma suggests a change in the tissue dynamic present during post-natal development. These differences highlight the importance of topographical information as to cell location during health and disease (51). Key open questions remain as to whether these clusters are driven by the immune response to allergens and if they are maintained during progression to adulthood. As with all early life studies in human, we are limited here by the lack of healthy controls and restricted to disease control subjects (56). However, although the pathway analysis implies some difference exist, this finding requires further analysis since the genes present in these pathways are from publicly available gene lists and thus may not provide a totally accurate reflection. In the future, additional spatial single cell RNAseq studies will be necessary to validate these findings.

In summary, our study demonstrates that tissue remodelling, induced by inhaled allergen, impacts blood vessel organisation during early life with a reduction of the pulmonary microcirculation that precedes other structural changes. This loss of blood vessels in specific areas may negatively impact gas exchange. Together, the need for increased oxygen supply due to tissue expansion and the concurrent reduction in pulmonary vasculature will lead to a quicker deterioration of lung and vascular function overall. Moreover, our study increases our fundamental understanding of temporal and regional responses during lung pathologies. Finally, we uncovered a new axis of regulation for tissue remodelling during allergic inflammation involving MCs and pericytes that requires further exploration during respiratory disorders (Figure 7).

**Figure 7:**
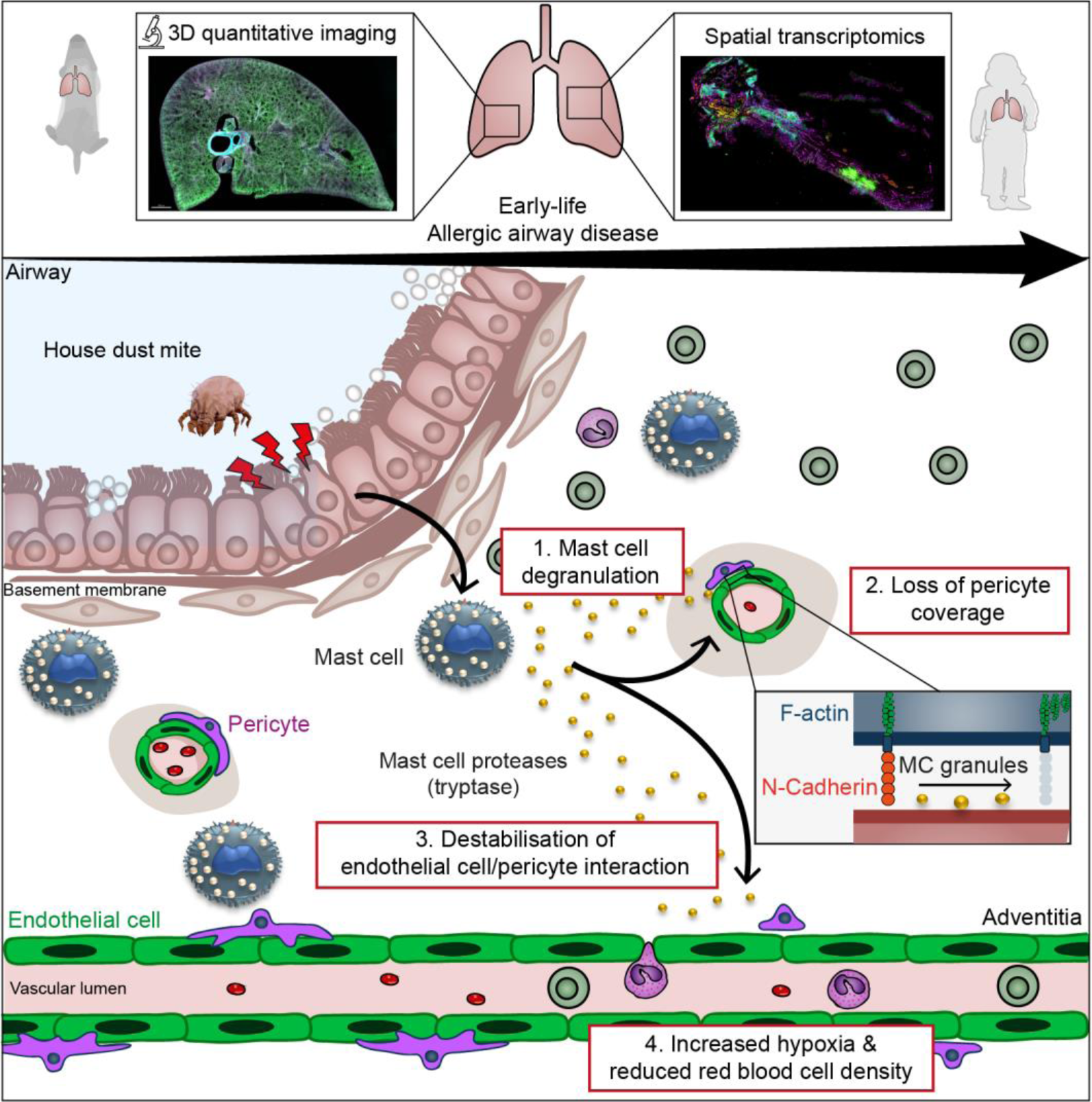
Graphical abstract. MC degranulation during early life allergic airway disease leads to pericyte damage and associated vascular remodelling in the lung adventitia.

## Methods

### Sex as a biological variable

Neonatal mice were of either sex, and litters were randomly assigned to control or experimental groups. Similar findings are reported for both sexes.

### Antibodies

Anti-mouse CD140b (Cat#136002, RRID: AB_1953332), APC anti-mouse CD140b antibody (Cat#136008 RRID: AB_2268091), Alexa Fluor® 647 anti-human CD31 Antibody (Cat#303111 RRID:AB_493077), Alexa Fluor® 488 anti-human CD31 Antibody (Cat#303109 RRID:AB_493075), APC anti-human CD140b (PDGFRβ) Antibody (Cat#323608 RRID:AB_2162787), Brilliant Violet 421™ anti-mouse CD45 Antibody (Cat# 103133 RRID:AB_10001045), BV421 anti-human CD117(C-kit) antibody (Cat# 313215 RRID:AB_10896056), BV421 anti-mouse CD117 (Cat# 560557 RRID:AB_1645258), PE/Cy5 anti-human CD45 (Cat# 304010 RRID:AB_314398), APC anti-mouse FCεRIα (Cat# 134316 RRID:AB_10640121), Brilliant Violet 711™ anti-human TNF-α Antibody (Cat# 502940 RRID AB_2563885), APC anti-mouse TER-119/Erythroid Cells antibody (Cat# 116212 RRID AB_313713), from Biolegend. Mouse/Rat CD31/PECAM-1 Antibody (Cat# AF3628 RRID:AB_2161028), Mouse Mast Cell Protease-6/Mcpt6 Antibody (Cat#MAB3736 RRID:AB_2240825), BV510 anti-human FCεRIα (Cat# 334626 RRID:AB_2564291), from R&D systems

### Animals

WT BALB/c (stock number #000651) were initially obtained from Charles River (UK) and maintained by in-house breeding. Each mother with its litter was housed separately. Mice were maintained in specific pathogen–free conditions and given food and water ad libitum. In individual experiments, all mice were matched for age and background strain.

### Study approvals

All in vivo experiments were conducted at the NHLI, Imperial College London, UK under the UK legislation for animal experimentation (PPL P996A24E1 and PP8328343) and in agreement with the UK Home Office Animals Scientific Procedures Act 1986 (ASPA).

Randomization: Neonatal mice were randomly assigned to control or experimental groups.

Sample size: The number of mice analysed for each different experimental approach is indicated on each figure.

### Human donors

Human adult samples used in this research project were obtained from the Imperial College Healthcare Tissue Bank (ICHTB). ICHTB is supported by the National Institute for Health Research (NIHR) Biomedical Research Centre based at Imperial College Healthcare NHS Trust and Imperial College London. ICHTB is approved by Wales REC3 to release human material for research (17/WA/0161), and the samples for this project (R22006) were issued from sub-collection reference number ICB_NC_21_017. The views expressed are those of the author(s) and not necessarily those of the NHS, the NIHR or the Department of Health.

School aged children (aged 8-17) that were undergoing a clinically indicated bronchoscopy, were recruited. Donor clinical data are available in Supplemental Table 1. The study was approved by the institutional ethics committee, and written, informed parental consent and child assent were obtained.

### Neonatal AAD

Neonatal mice were exposed repeatedly to either HDM (Greer Lot#360923) or PBS, intranasally from PN d. During the first 2 weeks of life, mice were administered 10 µg of HDM extract in 10 µl of PBS, three times a week. From week 3, mice received 15 µg of HDM in 15 µl of PBS. All outputs were assessed at 24 hours after allergen challenge (57).

### Precision-cut lung slices (PCLS)

PCLSs provided a 3D cell culture model to image vascular remodelling within the lung microenvironment and was adapted from previously described protocols (22, 58).

For mouse PLCS, lungs were inflated *in situ* with 0.4 ml of 2% low-melting agarose (Thermo Fisher Scientific) in PBS. After inflation, lungs were carefully dissected out and fixed in 4% Paraformaldehyde (PFA, Electron Microscopy Sciences) overnight at 4° C. Human lung tissue was collected from anonymised donors at Hammersmith and Royal Brompton hospitals NHS trusts and immediately fixed in 4% PFA overnight at 4°C. Following fixation, 100-200 µm transverse sections were prepared using a Compresstome VF-300 vibrating microtome (Precisionary Instruments).

PCLS were permeabilised in PBS complemented with 0.5% Triton (Sigma) for 1 hour at room temperature then blocked in animal-free blocker (2BSCIENTIFIC LTD) for 1 hour. Slices were incubated with indicated primary antibodies overnight at 4° C in 25% animal free blocker in PBS, and where required PCLS were incubated with secondary antibodies for 5 hours at room temperature in 25% animal free blocker in PBS. Lungs slices were mounted on microscopic slides (ThermoFisher) and immerged in ProLong Diamond (ThermoFisher) and kept at 4° C until image acquisition.

### Image acquisition

Images were acquired on a LEICA SP4 or SP8 (LEICA) using 20X objective (NA 0.7 and 0.75 for SP4 and SP8 respectively) or 10X objective (NA 0.4 for SP4 and SP8) with a resolution of 512x512 or 1024x1024 pixels. Motorised stages of the microscopes were employed for tile scan imaging and merged using LEICA built-in software (LAS) with a 10% overlap threshold. Adventitial regions (bronchovascular region) were defined as a lung region with the presence of a large airway (i.e. distinct stratified epithelium) and intermediate/large blood vessel. Vasculature associated to adventitial region was analysed within a 150 µm radius from the main airway/large vessel. Parenchymal regions were identified morphologically based on the alveoli structure more than 300 µm from any large airways.

### Image analysis

Image analysis and rendering were performed using IMARIS 8.1 or 9.3 (Bitplane). LIF files were converted into IMARIS .ims files using IMARIS Converter software v9.9.1 (Bitplane). Cell, HIF-1α, avidin^+^ granules, m-MCP6^+^ spot number was analysed using the semi-automatic spot function (cell diameter 10 and 15 µm for leukocytes and pericytes respectively, spot size 1-5 µm for vesicles and granules) and volumes (i.e. EC/pericyte coverage) were determined using the surface function. Numbers of cells were normalised to the total volume of the image. Thresholds for cell volume analysis in PCLS sections were maintained across the experimental conditions to provide accurate comparisons. Pericyte and endothelial cell volumes were determined using the cell function of IMARIS and similar thresholds for cell cytoplasm were used across all the experimental conditions using either CMTMR staining or F-actin. All image analysis was performed on raw images without fluorescence modification.

### Cell purification

Lung tissues were gently dissociated in small fragments (1-5 mm length) using scissors then placed in 6 wells plates (Corning). To generate pericytes, mouse and human lung samples were differentiated as previously described with small modifications (59). Briefly, cells were grown in Dulbecco’s Modified Eagle’s Medium (DMEM) supplemented with 10% FCS, 100 U/mL penicillin, 100 mg/mL streptomycin (all Thermo Fisher Scientific) and 100 pM pigment epithelium-derived factor (PEDF, Sigma-Aldrich). After 15 to 22 days of culture, cells were detached using trypsin (Thermo Fisher) and isolated using the EasySep™ APC Positive Selection Kit II (StemCell) according to manufacturer instructions. APC conjugated anti-PDGFRβ was used for magnetic cell sorting. Cell purity (PDGFRβ^+^NG2^-^) was assessed by flow cytometry. Of note, human pericytes were smaller compared to mouse pericytes.

Mouse lung MCs were grown in OPTI-MEM supplemented with 10% FCS, 100 U/mL penicillin, 100 mg/mL streptomycin and 4% CHO transfectants secreting murine Stem Cell Factor (SCF, a gift from Dr P. Dubreuil, Marseille, France, 4% correspond to ∼50 ng/ml SCF) for 6-8 weeks. Human lung MCs were grown in OPTI-MEM supplemented with 2.5% BSA, 100 U/mL penicillin, 100 mg/mL streptomycin and 4% CHO transfectants secreting murine SCF for ∼8 weeks. Mast cell purity was assessed by flow cytometry (CD117^+^/FcεRI^+^).

### Functional assays

Mouse or human pericytes (∼10,000 cells) were seeded onto µ-Slide 8 Well high ibiTreat: tissue culture treated (Ibidi) and leave to rest for 24 hours. Cells were stained using CellTracker™ Orange CMTMR Dye (ThermoFisher) according to manufacture instructions. ∼5,000 previously anti-dinitrophenyl (DNP) IgE-sensitized lung MCs (overnight at 1 µg/ml, clone SPE-7, Merck) were washed in Tyrode buffer (Merck) and placed on top of pericytes for ∼15 minutes at 37° C. Increasing concentrations of DNP-BSA (Merck) were added and cells were left for 24 hours at 37° C. In some experiments, protease inhibitor cocktail (Merck) was used according to manufacturer instructions or pericytes were directly exposed to recombinant m-MCP6 (R&D systems, matured according to manufacturer instructions) or 10 µM of APC-366 (tryptase inhibitor, Merck) was employed to block human tryptase. Following incubation, cells were placed on ice and stained for surface markers for 30 minutes, washed and fixed for ∼15 minutes with 4% PFA at 37° C. Where necessary, fixed cells were permeabilised using Permeabilization Buffer (ThermoFisher) and stained for 30 minutes at room temperature with fluorochrome-conjugated antibodies or for F-actin (Alexa Fluor™ 488 Phalloidin, ThermoFisher). Cells were kept in PBS before image acquisition. Mast cell degranulation flow cytometry assay were performed as previously described (31). Briefly, ∼10000 IgE sensitized lung MCs were washed in Tyrode buffer (Merck) and placed in a 96 wells U bottom plates at 37°C. Cells were stimulated with increasing DNP-BSA concentrations (Merck) for 30 minutes at 37°C, collected and processed for flow cytometry.

### Flow cytometry

Cells were resuspended and washed in PBS + 0.5% BSA + 0.1 mM EDTA. Primary antibodies were incubated for ∼30 minutes at 4° C, washed 2 times and incubated with secondary antibodies if necessary for another 30 minutes at 4° C. Data were acquired on a BD LSR Fortessa using FACSDIVA software (both from BD Biosciences) and analysed using FlowJo software (v10, FlowJo).

### scRNAseq analysis

Mouse and human pericyte marker analysis was performed using the publicly available data mining website (http://lungendothelialcellatlas.com/) (24). Data were visualised using the multi-gene query options and the expression of the best 8 markers for pericytes were analysed in mouse and human scRNAseq datasets (30000 cells combined from 5 cohorts).

### Spatial transcriptomics

We followed experimental methods (NanoString) described in the publication (37). Briefly, FFPE human lung samples were baked overnight at 37° C followed by 3 hours of baking at 65° C, then loaded onto a Leica Bond RX Fully Automated Research Stainer for subsequent processing steps. The processing protocol included three major steps: 1) slide baking, 2) antigen retrieval for 20 min at 100° C, and 3) treatment with Proteinase K (1.0 µg/mL in 1xPBS) for 15 min. Following these steps, slides were removed from the Leica Bond RX, and a cocktail of GeoMx Cancer Transcriptome Atlas probes (specific to ∼1,800 genes separate targets) were applied to each slide and allowed to hybridize at 37° C overnight in a humidity chamber. The following day, slides were washed, blocked, and allowed to incubate with a combination of Alexa Fluor 488-labeled anti-alpha smooth muscle actin antibody (Invitrogen/Thermo product #: 53-9760-82; clone: 1A4), Alexa Fluor 594-labeled Anti-vimentin antibody (Santa Cruz product #: sc-373717 AF594; clone: E-5), Alexa Fluor 647-labeled anti-CD45 antibody (Cell Signaling Technology product #: 13917BF; clone: D9M8I), and Syto83 nucleic acid stain. Slides were stained for 1 h at room temperature in a humidity chamber. Slides were then washed and loaded onto a GeoMx instrument. On the GeoMx machine, slides were fluorescently scanned, and ROIs were collected from the following areas: smooth muscle, epithelium, fibroblast (vessel-rich), and immune-rich. The GeoMx device exposed ROIs to 385 nm light (UV), releasing the indexing oligos. Indexing oligos were collected with a microcapillary and deposited into a 96-well plate. Samples were dried down overnight, then resuspended in 10 µL of DEPC-treated water. PCR was performed using 4 µL of each sample and the oligos from each ROI were indexed using unique i5 and i7 dual-indexing systems (Illumina). PCR reactions were purified twice using AMPure XP beads (Beckman Coulter, Inc) according to the manufacturer’s protocol. Purified libraries were sequenced on an Illumina NovaSeq 6000. Data analysis was performed as described (37). Following removal of targets consistently below the limit of quantification (i.e. <5000 raw reads) and negative probes, the limit of detection above which a gene was called “detected” was defined as 2 standard deviations above the geometric mean of negative probes. Datasets were normalized using upper quartile (Q3) normalization. Data analysis was then performed using the DSP platform and R software. Cell deconvolution analysis was performed using the following R packages: GSVA, Stringr and Dplyr (60). Original single cell RNAseq data for cell deconvolution were obtained from Deprez et al. (38).

### Statistical analysis

Most statistical analyses including PCA were performed using Prism software (GraphPad v9.4.1). Pathway analysis (GSEA) analysis was performed on the DSP platform using a Mann-Whitney test corrected for multiple testing using Benjamini-Hochberg (BH) procedure and data were represented using Prism software. t-SNE analysis was performed using build-in functions of FlowJo software, iterations were fixed at 1000, perplexity 30 and learning rate 22. The results are expressed as means ± SEM and the n numbers for each dataset are provided in the Figure legends. Comparisons between two groups were carried out using the paired or unpaired Student’s or Mann-Whitney t test as appropriate. One-way ANOVA followed by Tukey post hoc test was performed for multiple group comparisons. Statistical significance was accepted at p < 0.05.

## Resource availability

Further information and requests for resources and reagents should be directed to and will be fulfilled by lead author Régis Joulia

## Data and Code availability

All data represented in this study are present in the Supporting data values file available. Spatial transcriptomic data are available as supplementary files attached to this paper. Any additional information required to reanalyse the data reported such as original source images in this paper is available from the lead contact upon request. The paper does not report original code.

## Author contributions

Conceptualization, R.J., F.P., C.M.L. Methodology, R.J., F.P. Investigation, R.J., F.P., H.S., A.V., M.G.M., L.J.E., Resources, W.T. M.A., K.B., E.S., P.M., R.J.H., S.A.W., L.Y., S.S. Formal analysis, R.J., F.P. and A.V. Visualization, R.J. Writing – original draft. R.J. Writing – review & editing, R.J., F.P., H.S., W.T., A.V., L.J.E., M.G.M., M.A., K.B., E.S., P.M., R.J.H., S.A.W., L.Y., S.S. and C.M.L. Supervision, R.J. and C.M.L.

## Acknowledgments

This work was supported by funds from the Wellcome Trust (107059/Z/15/Z and 220254/Z/20/Z) to C.M.L. R.J. is supported by fellowships from the British Heart Foundation Imperial College Centre for Research Excellence (RE/18/4/34215) and ACTERIA EFIS Foundation Allergology 2023. The Facility for Imaging by Light Microscopy (FILM) at Imperial College London is part-supported by funding from the Wellcome Trust (grant 104931/Z/14/Z) and the authors would like to thank Dr. David Gaboriau for invaluable help. This project was supported in part by the NIHR Imperial Biomedical Research Centre (BRC). The views expressed are those of the authors and not necessarily those of the NIHR or the Department of Health and Social Care.

