## Supplemental material for "Mast cell activation disrupts interactions between endothelial cells and pericytes during early life allergic asthma"

**Supplemental Information**

Joulia et al.

- **Supplemental Figure 1.** PDGFRβ is reliable marker for murine and human lung pericytes.
- **Supplemental Figure 2.** Parameters of vascular remodelling in neonate mice exposed with HDM.
- **Supplemental Figure 3.** HDM rechallenge led to loss of pericyte coverage and MC activation.
- **Supplemental Figure 4.** Early life allergen exposure does not lead to immune cell infiltration in the lung parenchyma.
- **Supplemental Figure 5.** Validation of primary mouse and human pericytes and MCs.
- **Supplemental Figure 6.** Pericyte and endothelial cell spatial transcriptomic profiles in children with asthma.

- **Supplementary Video 1 (related to Fig. 1).** PCLS of the lung vasculature during early life.
- **Supplementary Video 2 (related to Fig. 2).** Loss of adventitial pericyte protrusions following HDM exposure.
- **Supplementary Video 3 (related to Fig. 2).** Distribution of CTMCs in the lung adventitia and analysis of MC degranulation.

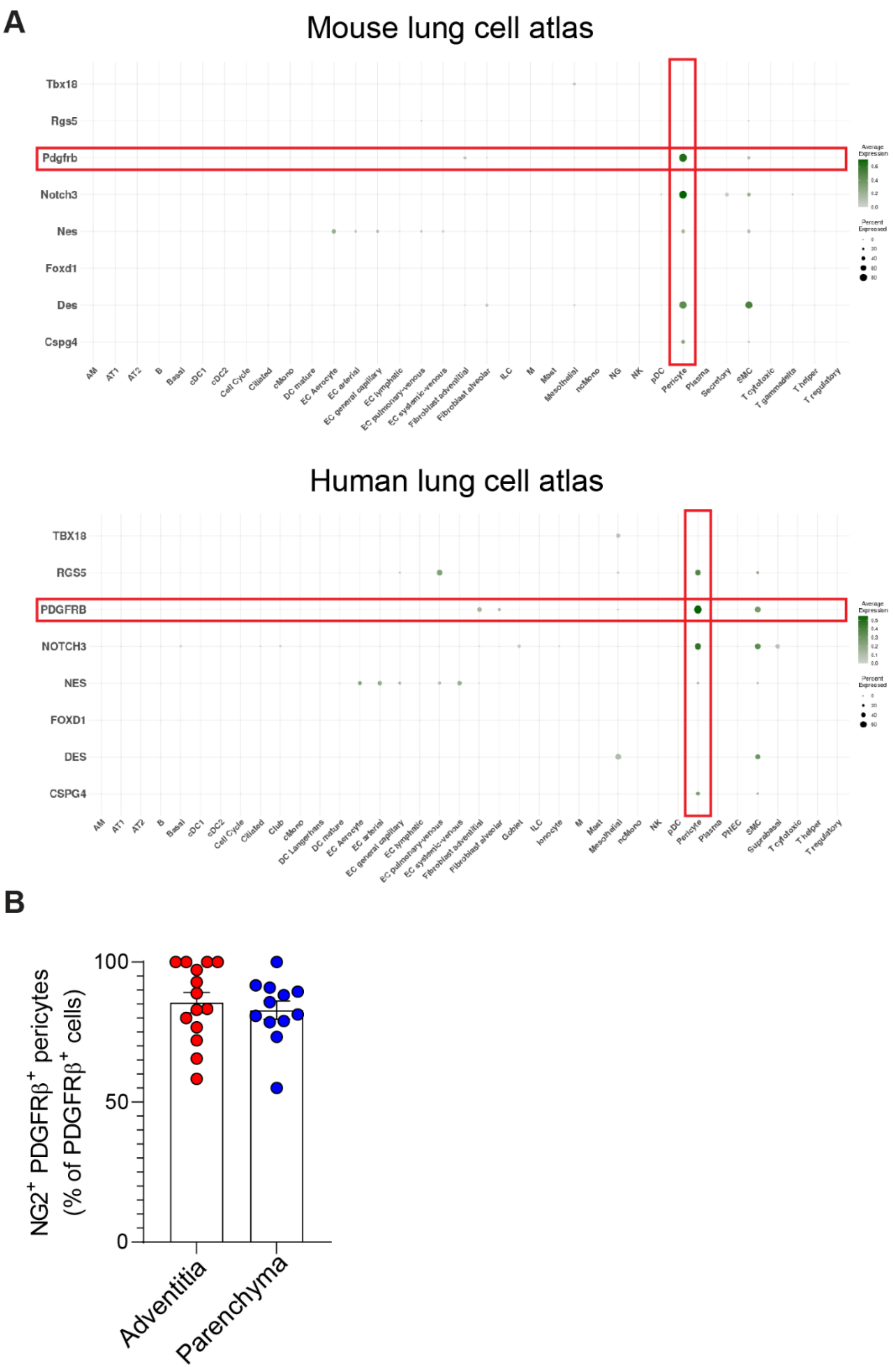


**Supplemental Figure 1. PDGFRβ$^{+}$ is reliable marker for murine and human lung pericytes.** (**A**) RNAseq data from the mouse and human lung cell atlas analysing the level of expression of 8 markers for pericytes in the lungs. Red boxes highlighted *Pdgfrb* gene expression. Average expression is colour coded and the size of the circle reflected the frequency of positive cells. (**B**) Colocalization analysis between PDGFRβ$^{+}$ and NG2$^{+}$ cells in the microcirculation of the lung adventitia and parenchyma (n=12/14 images from 3 mice). Mean ± SEM.

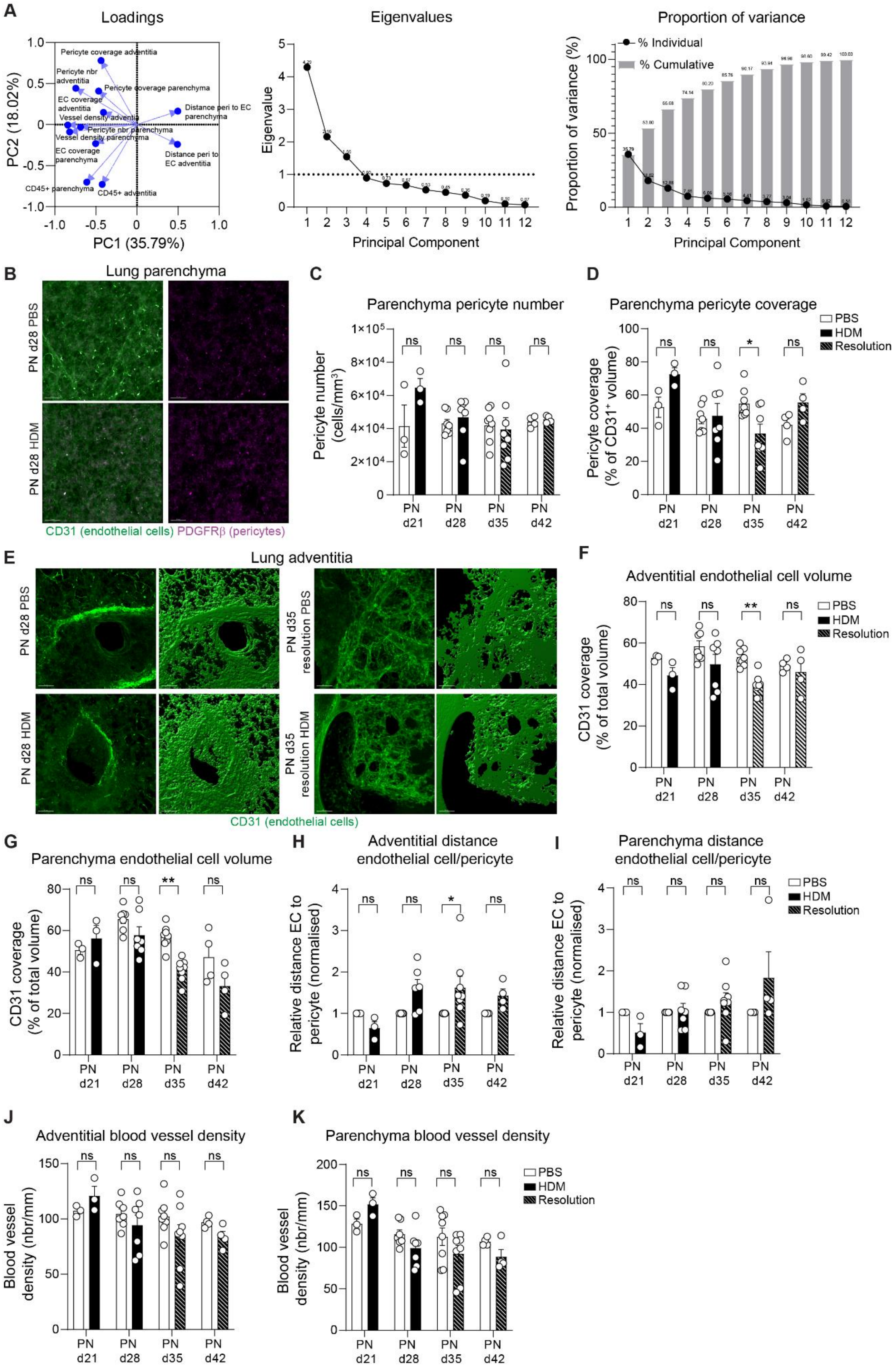
A
Loadings
Pericyte coverage adventitia
Pericyte nbr adventitia
Pericyte coverage parenchyma
EC coverage adventitia
Vessel density adventia
Distance peri to EC parenchyma
Pericyte nbr parenchyma
Vessel density parenchyma
EC coverage parenchyma
Distance peri to EC adventitia
CD45+ parenchyma
CD45+ adventitia
PC2 (18.02%)
PC1 (35.79%)
Eigenvalues
Eigenvalue
Principal Component
Proportion of variance
% Individual
% Cumulative
Proportion of variance (%)
Principal Component
B
Lung parenchyma
PN d28 PBS
PN d28 HDM
CD31 (endothelial cells) PDGFRβ (pericytes)
C
Parenchyma pericyte number
Pericyte number (cells/mm3)
D
Parenchyma pericyte coverage
Pericyte coverage (% of CD31+ volume)
PBS
HDM
Resolution
E
Lung adventitia
PN d28 PBS
PN d28 HDM
PN d35 resolution PBS
PN d35 resolution HDM
CD31 (endothelial cells)
F
Adventitial endothelial cell volume
CD31 coverage (% of total volume)
G
Parenchyma endothelial cell volume
CD31 coverage (% of total volume)
H
Adventitial distance endothelial cell/pericyte
Relative distance EC to pericyte (normalised)
I
Parenchyma distance endothelial cell/pericyte
Relative distance EC to pericyte (normalised)
J
Adventitial blood vessel density
Blood vessel density (nbr/mm)
K
Parenchyma blood vessel density
Blood vessel density (nbr/mm)
PN d21
PN d28
PN d35
PN d42

**Supplemental Figure 2. Parameters of vascular remodelling in neonate mice exposed with HDM.** Neonate mice were exposed with PBS or HDM as indicated in Figure 1D. (**A**) PCA loading values, length of the vectors are proportional of their importance in the principal components, eigenvalues and proportion of variance for the PCA analysis presented in Figure 1F. (**B**) 3D rendering of a PCLS section in PBS and HDM exposed mice 3 weeks post first inhalation showing CD31 (green, endothelial cells) and PDGFRβ (magenta, pericytes), scale bars 50 µm (representative of 3 independent experiments). (**C-D**) PDGFRβ$^{+}$ pericyte number (**C**) and coverage (**D**, normalised to the total volume of CD31$^{+}$ blood vessel) in the lung parenchyma (n=3-8 mice per group from 4 independent experiments). (**E**) 3D rendering of a PCLS section in PBS and HDM exposed mice 3 weeks (left panels) and left to rest for 1 week showing CD31 (green, endothelial cells). (**F-G**) Adventitial (**F**) and parenchyma (**G**) endothelial cell coverage (normalised to the total volume of the image, n=3-8 mice per group from 4 independent experiments). (**H-I**) Distance between endothelial cell and pericyte (normalised to PBS exposed mice) in the lung adventitia (**H**) and parenchyma (**I**, n=3-8 mice per group from 4 independent experiments). (**J-K**) Blood vessel density in the lung adventitia (**J**) and parenchyma (**K**, n=3-8 mice per group from 4 independent experiments). Mean ± SEM. two-way ANOVA followed by Sidak's post-hoc test, *$p<0.05$, **$p<0.01$,***$p<0.001$, ns=not significant.

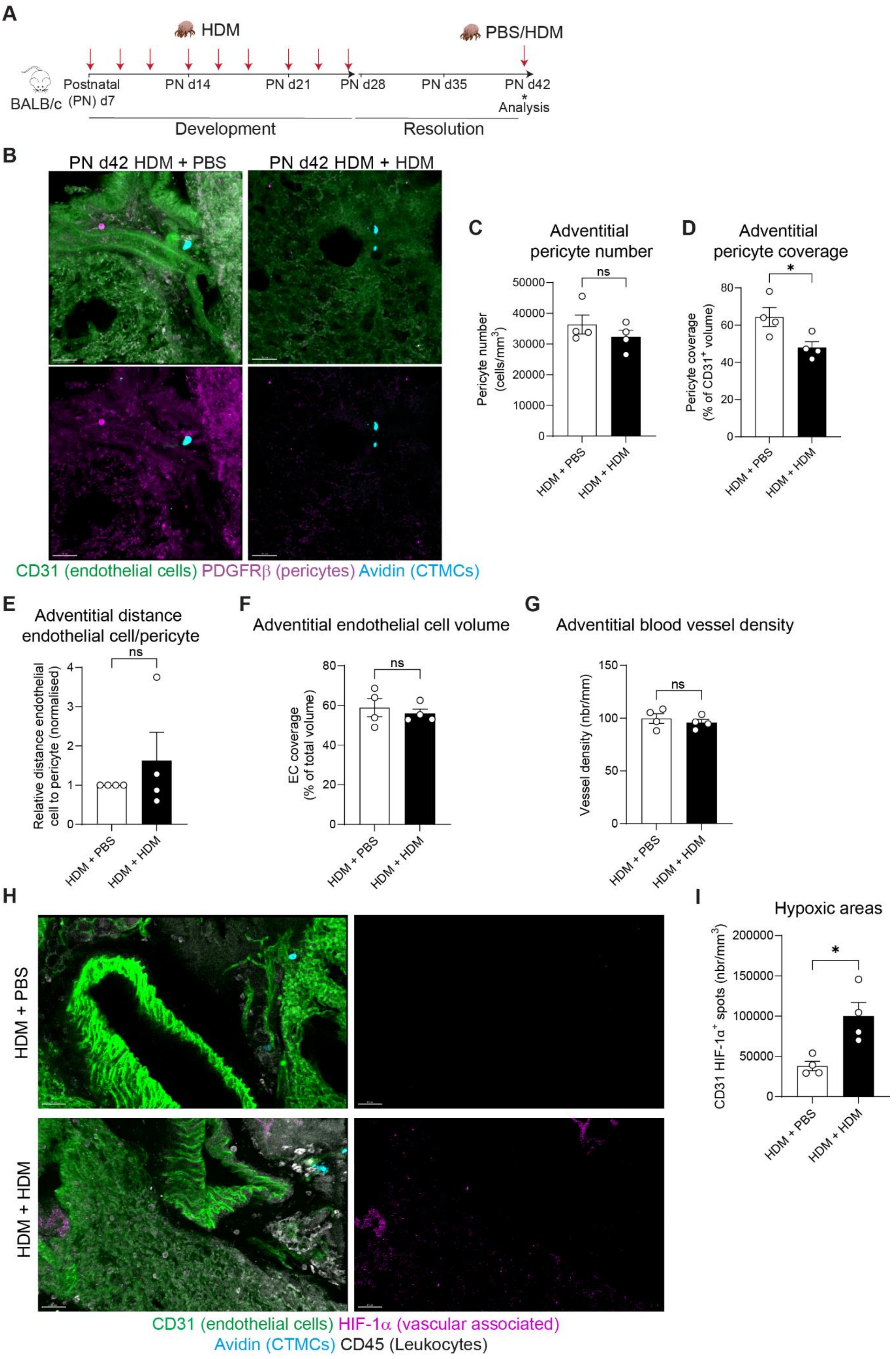
A
HDM
PBS/HDM
BALB/c
Postnatal (PN) d7
PN d14
PN d21
PN d28
PN d35
PN d42
* Analysis
Development
Resolution
B
PN d42 HDM + PBS
PN d42 HDM + HDM
CD31 (endothelial cells) PDGFRβ (pericytes) Avidin (CTMCs)
C
Adventitial pericyte number
Pericyte number (cells/mm3)
ns
HDM + PBS
HDM + HDM
D
Adventitial pericyte coverage
Pericyte coverage (% of CD31+ volume)
*
E
Adventitial distance endothelial cell/pericyte
Relative distance endothelial cell to pericyte (normalised)
F
Adventitial endothelial cell volume
EC coverage (% of total volume)
G
Adventitial blood vessel density
Vessel density (nbr/mm)
H
HDM + PBS
HDM + HDM
CD31 (endothelial cells) HIF-1α (vascular associated)
Avidin (CTMCs) CD45 (Leukocytes)
I
Hypoxic areas
CD31 HIF-1α+ spots (nbr/mm3)

**Supplemental Figure 3. HDM rechallenge led to loss of pericyte coverage and MC activation.** (**A**) BALB/c mice aged 7 days were exposed to intermittent intranasal HDM for 3 weeks (red arrows), following 2 weeks of rest mice were rechallenged with a single dose of HDM or PBS. (B) PCLS of PBS or HDM exposed mice (PN d42) showing CD31 (green, endothelial cells), PDGFRβ (magenta, pericytes) and avidin (cyan, CTMCs) in an adventitial region, scale bars 50 µm. (**C-D**) PDGFRβ$^{+}$ pericyte number (**C**) and coverage (**D**) (normalised to the total volume of CD31$^{+}$ blood vessel) in the lung adventitia (n=4 mice per group). (**E**) Distance between endothelial cell and pericyte (normalised to PBS exposed mice) in the lung adventitia (n=4 mice per group). (**F**) Adventitial endothelial cell coverage (normalised to the total volume of the image, n=4 mice per group). (**G**) Blood vessel density in the lung adventitia (n=4 mice per group). (**H**) Representative PCLS of PBS or HDM exposed mice (PN d42) showing CD31 (green, endothelial cells), HIF-1α (magenta), avidin (cyan, CTMCs) and CD45 (white, leukocytes) in an adventitial region, scale bars 40 µm. (**I**) Number of HIF-1α spots in adventitial region in PBS and HDM exposed mice (n= 4 mice per group). Mean ± SEM, two tailed Student t-test, *$p<0.05$, ns=not significant.

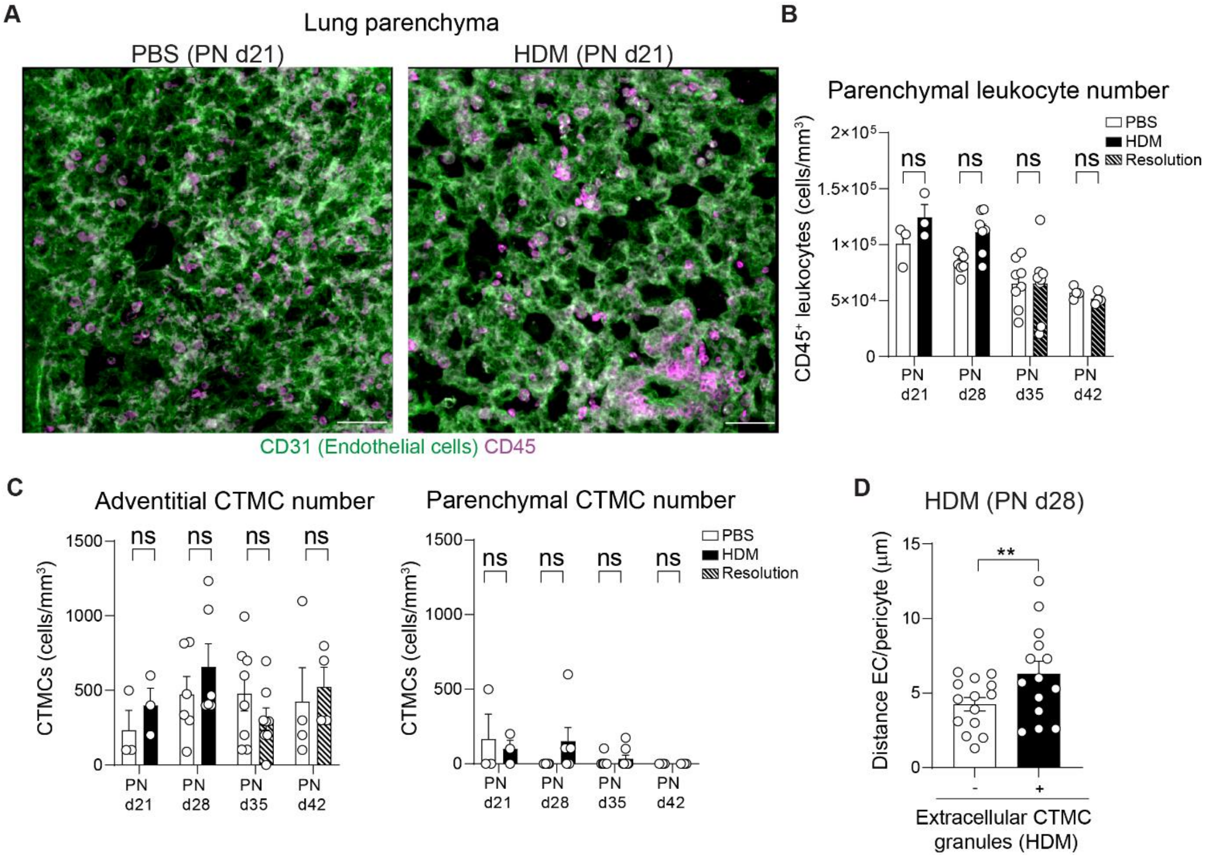


**Supplemental Figure 4. Early life allergen exposure does not lead to immune cell infiltration in the lung parenchyma.** Neonate mice were exposed to PBS or HDM as indicated in Fig. 1D. (**A**) 3D rendering of a PCLS section in the lung parenchyma in PBS and HDM exposed mice 3 weeks post beginning of the treatment showing CD31 (green, ECs) and CD45 (magenta, leukocytes), scale bars 50 µm (representative from 4 independent experiments). (**B**) $CD45^+$ number (n=3-8 mice per group from 4 independent experiments). (**C**) CTMC number in the lung adventitia (left panel) and parenchyma (right) number (n=3-8 mice per group from 4 independent experiments). (**D**) Distance between endothelial cells and pericytes at site of MC degranulation (less than 40 µm of MC granules or control regions without MC granules; n=14 images from 4 HDM treated mice). Mean ± SEM. **B, C**, two-way ANOVA followed by Sidak's post-hoc test; **D**, two-tailed Student's t-test. **p<0.01, ns=not significant.

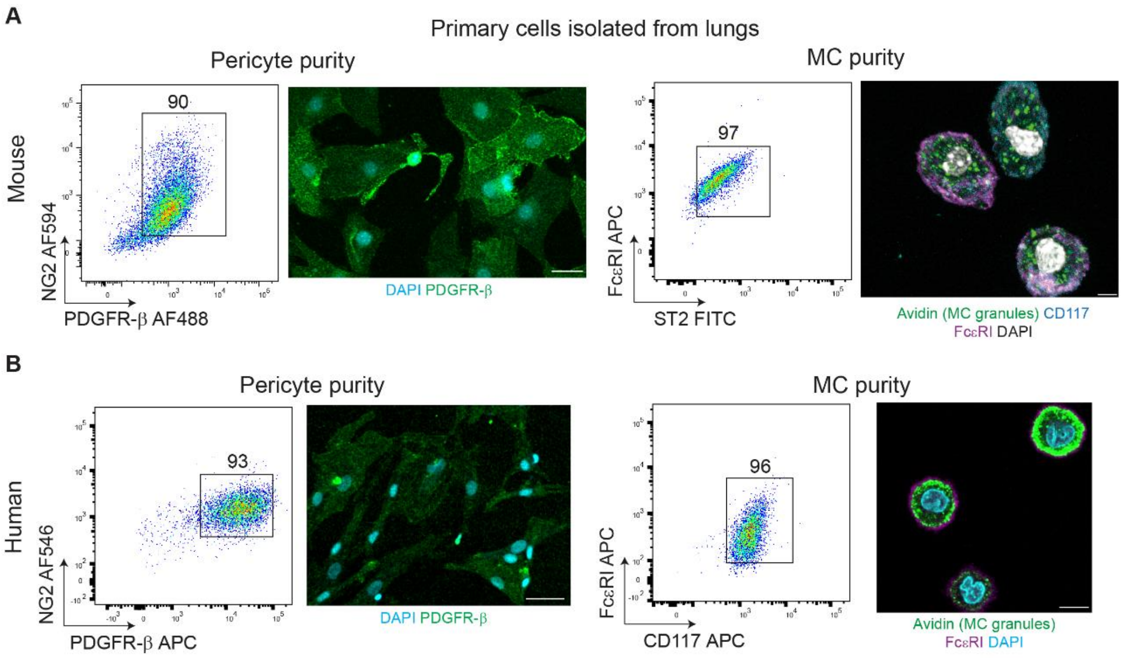


**Supplemental Figure 5. Validation of primary mouse and human pericytes and MCs.** Mouse and human cells were isolated from lung parenchyma and differentiated in their respective media (see methods). (**A**) Flow cytometry profiles and confocal images of mouse pericytes and MCs showing a high purity level for pericytes (i.e. $NG2^+/PDGFR\beta^+$) and MCs ($Fc\varepsilon RI^+/ST2^+/CD117^+$), scale bars 50 µm (pericytes) and 5 µm (MCs) (representative 4 independent experiments). (**B**) Flow cytometry profiles and confocal images of human pericytes and MCs showing high purity level for pericytes (i.e. $NG2^+/PDGFR\beta^+$) and MCs ($Fc\varepsilon RI^+/CD117^+$), scale bars 50 µm (pericytes) and 10 µm (MCs) (representative 4 independent experiments).

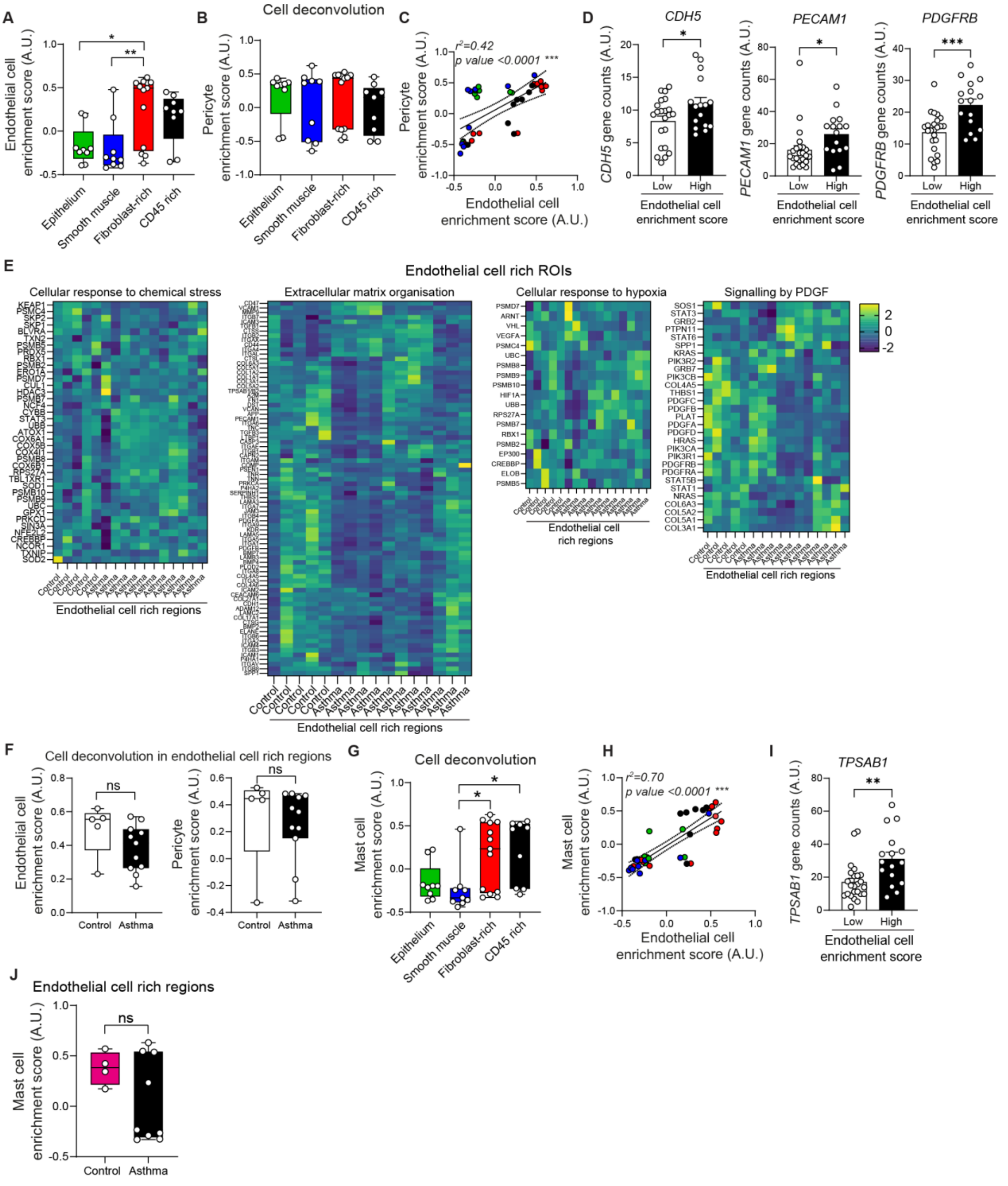


**Supplemental Figure 6. Pericyte, endothelial cell and MC spatial transcriptomic profiles in children with asthma.** Human endobronchial biopsies were processed for spatial transcriptomics using the Nanostring GeoMx® platform and analysed using the Cancer Transcriptome Atlas (CTA, ~1,800 genes). (**A**) Endothelial cell enrichment score (n=9-13 ROIs per group from 2 controls and 4 asthma). (**B**) Pericyte enrichment score (n=9-13 ROIs per group from 2 controls and 4 asthma). (**C**) Correlation between endothelial cell and pericyte enrichment score (n=40 ROIs from 2 controls and 4 asthma, green epithelium, red fibroblast rich, blue smooth muscle, black immune rich). (**D**) Gene counts for indicated genes in endothelial rich regions and rest of ROIs (n=16-24 ROIs from 2 controls and 4 asthma). (**E**) Gene expression of indicated pathway from control and asthma endothelial cell rich ROIs represented as heatmap. (**F**)

Endothelial cell and pericyte enrichment score in endothelial cell rich regions (n=5-11 ROIs per group from 2 controls and 2 asthma). (**G**) MC enrichment score (n=9-13 ROIs per group from 2 controls and 4 asthma). (**H**) Correlation between MC and endothelial cell enrichment score (n=40 ROIs from 2 controls and 4 asthma, green epithelium, red fibroblast rich, blue smooth muscle, black immune rich). (**I**) Gene counts for *TPSAB1* in endothelial rich regions and rest of ROIs (n=16-24 ROIs). (**J**) MC enrichment score in endothelial cell rich regions (n=4-9 ROIs per group from 2 controls and 2 asthma). **A**, B, G one-way ANOVA followed by Tukey's post-hoc test, **C**, **D**, **F, I, J** two-tailed Student's t-test. *p<0.05, **p<0.01, ***p<0.001, ns= not significant.

**Supplemental video legends:**

**Supplemental Video 1 (related to Figure 1): PCLS of the lung vasculature during early life.** Mouse WT lungs (aged 4 weeks) were inflated with 2% agarose, isolated and swiftly fixed in 4% PFA overnight. PCLS (200 µm thickness) were stained for CD31 (EC, green), α-SMA (SMC, cyan) and PDGFRβ (pericyte, purple). Tile-scan imaging was performed using a 10X objective and reconstructed using IMARIS software. The video shows the parenchyma and adventitial vasculature network with the adventitia regions highlighted.

**Supplemental Video 2 (related to Figure 2): Loss of adventitial pericyte protrusions following HDM exposure.** Neonate mice were exposed with PBS or HDM as indicated in Fig. 1D. Lungs were collected 24 hours following the last dose of HDM and processed as indicated in Video 1 legend. Zoom in images of lung adventitia show the endothelial cell and pericyte network next to a large airway and large blood vessel in PBS and HDM exposed mice. IMARIS software was used to model pericyte cell body (red dots) and cell protrusion (purple surface). The movie illustrates how pericyte cell protrusions are lost following exposure to 3 weeks of HDM.

**Supplemental Video 3 (related to Figure 2): Distribution of CTMCs in the lung adventitia and analysis of MC degranulation.** Neonate mice were exposed with PBS or HDM as indicated in Fig. 1D. Lungs were collected 24 hours following the last dose of HDM and processed as indicated in Video 1 legend. PCLS were stained for avidin (MC granules, blue) and blood vessels (CD31, green). Low magnification image shows the distribution of MCs around a large airway and adjacent to large blood vessels. Zoom in image shows the degranulated profile of multiple CTMCs and the IMARIS analysis of MC granule volume (colour coded according to the volume).
